# Deciphering the Biosynthetic Potential of Microbial Genomes Using a BGC Language Processing Neural Network Model

**DOI:** 10.1101/2023.11.30.569352

**Authors:** Qilong Lai, Shuai Yao, Yuguo Zha, Haobo Zhang, Ying Ye, Yonghui Zhang, Hong Bai, Kang Ning

## Abstract

Microbial secondary metabolites are usually synthesized by colocalized genes termed biosynthetic gene clusters (BGCs). A large portion of BGCs remain undiscovered in microbial genomes and metagenomes, representing a pressing challenge in unlocking the full potential of natural product diversity. In this work, we propose BGC-Prophet, a language model based on the transformer encoder that captures the distant location-dependent relationships among biosynthetic genes, allows accurately and efficiently identifies known BGCs and extrapolates novel BGCs among the microbial universe. BGC-Prophet is the first ultrahigh-throughput (UHT) method that is several orders of magnitude faster than existing tools such as DeepBGC, enabling pan-phylogenetic screening and whole-metagenome screening of BGCs. By analyzing 85,203 genomes and 9,428 metagenomes, new insights have been obtained about the diversity of BGCs on genomes from the majority of bacterial and archaeal lineages. The profound enrichment of BGCs in microbes after important geological events have been revealed: Both the Great Oxidation and Cambrian Explosion events led to a surge in BGC diversity and abundance, particularly in polyketides. These findings suggest that it is a general but constantly evolving approach for microbes to produce secondary metabolites for their adaptation in the changing environment. Taken together, BGC-Prophet enables accurate and fast detection of BGCs on a large scale, holds great promise for expanding BGC knowledge, and sheds light on the evolutionary patterns of BGCs for possible applications in synthetic biology.

**Highlights:**

- BGC-Prophet shows superior performance to existing tools in terms of accuracy and speed.
- BGC-Prophet is the first ultrahigh-throughput (UHT) method that enables pan-phylogenetic screening and whole-metagenome screening of BGCs.
- BGC-Prophet builds the comprehensive profile of BGCs on 85,203 genomes and 9,428 metagenomes from the majority of bacterial and archaeal lineages.
- BGC-Prophet reveals the profound enrichment pattern of BGCs after important geological events.

## Introduction

Microbial secondary metabolism, one of the important sources of natural products, is generated through the coordinated action of numerous genes organized into biosynthetic gene clusters (BGCs) [1, 2]. Across the tree of life, these natural products comprise thousands of different chemical structures, including polyketides, saccharides, terpenes and alkaloids, that facilitate an organism’s ability to thrive in a particular environment [3, 4]. These secondary metabolites also demonstrate efficacy across multiple therapeutic areas, including antimicrobial and cancer immunotherapy [5, 6]. The biosynthesis of these compounds involves multienzyme loci called BGCs, which encode the biosynthetic pathways for one or more specific compounds [7, 8]. With the exponential growth of genomic data, identifying and classifying BGCs from microbial genomes or metagenomic assembled genomes (MAGs) has become a pressing challenge in exploring and exploiting natural product diversity [9, 10]. Developments in computational omics technologies have provided new means to assess the hidden diversity of natural products, unearthing new potential for drug discovery [11, 12].

BGC encodes a series of genes involved in biosynthetic or metabolic pathways, which are arranged in a sequential order on the genome. These genes work together to produce one or more small molecular compounds, such as penicillin [13, 14]. Recent insights revealed that BGC comprised a cluster of spatially adjacent colocalization genes, including biosynthetic genes and auxiliary genes (*e.g.*, transport-related genes, regulatory genes) [15, 16]. These biosynthetic genes play key catalytic roles in the formation of microbial secondary metabolites. In addition to biosynthetic enzymes, many BGCs also harbor enzymes to synthesize specialized monomers for a pathway. For example, the erythromycin gene cluster encodes a set of enzymes for the biosynthesis of two deoxy-sugars that are appended to the polyketide aglycone [17]. In many cases, transporters, regulatory elements, and genes that mediate host resistance are also contained within the BGC [18]. Although some BGCs are so well understood that the biosynthesis of their small molecule product has been reconstituted in heterologous hosts, little is known about the vast majority of BGCs, even those that have been linked to a small molecule product.

The explosion of microbial genomic data, including complete and partial genome sequences, has led to a transformative change in how computational methods are employed in natural product drug candidate discovery. Computational approaches are being developed to predict BGCs based on genome sequences alone, fuelled by data on known biosynthetic pathways and their chemical products, which are currently standardized with predicted BGCs stored in public databases [19]. Identifying natural product BGCs still largely relies on rule-based methods such as those used in antiSMASH [15, 16] and PRISM [20]. Although these approaches are successful at detecting known BGC categories, they are less proficient at identifying novel categories of BGC [21, 22]. In these more complex cases of identifying novel BGCs, machine learning algorithms have been shown to offer significant advantages over rule-based methods. For example, ClusterFinder [23], NeuRiPP [24] and DeepRiPP [25] each use machine learning to identify BGCs. These methods often have a tradeoff in terms of efficiency and accuracy, have a higher false positive rate than rule-based approaches and suffer from false negatives for known categories of BGC. Recently, deep learning approaches have been developed for BGC annotation, including DeepBGC [26], e-DeepBGC [27], Deep-BGCPred [28], and SanntiS [29]. All of these deep learning approaches call biosynthetic gene families using collections of curated profile-Hidden Markov Models (pHMMs) and employs a bidirectional long short-term memory (BiLSTM) recurrent neural network for improved identification of BGCs [26-29]. Although these approaches have improved the detection of BGCs from bacterial genomes and harness great potential to detect novel categories of BGCs, they have common drawbacks: BiLSTM might lose distant memories during the recurrent neural network and is unable to capture distant location-dependent relationships between biosynthetic genes, while the utilization of pHMM heavily relies on manual determination by experts to define the scope of each domain from the Pfam database [30], and is computationally intensive.

Collectively, several challenges persist for contemporary BGC prediction tools. First, these tools cannot accurately capture the location-dependent relationships between genes, resulting in limited accuracy and applicability, particularly in novel BGC predictions. Additionally, existing methods rely on time-consuming sequence alignment to extract features (such as Pfam domains), which hinders their speed for pan-phylogenetic screening and whole-metagenome screening of BGCs. Furthermore, the low throughput of existing methods makes it impossible for them to construct a comprehensive profile of BGCs on almost all lineages of genomes and metagenomes, thereby precluding the revelation of enrichment patterns of BGCs on a broad scale.

To address these limitations, we proposed BGC-Prophet, a deep learning approach that leverages a language model to accurately and efficiently identify known BGCs and extrapolate novel BGCs among the microbial universe. Previous studies have shown that the success of language models for BGC detection and product classification [31, 32]. Encouraged by this, our BGC-Prophet employs the powerful language model of the transformer encoder [33, 34], which captures the distant location-dependent relationships among biosynthetic genes for improved BGC detection and classification.

Our experiments show that BGC-Prophet achieves a >90% area under the receiver operating characteristic curve (AUROC) on the validation datasets and offers a comparable ability in BGC identification to existing tools such as DeepBGC. BGC-Prophet is the first ultrahigh-throughput (UHT) method that is several orders of magnitude faster than existing tools such as DeepBGC, enabling pan-phylogenetic screening and whole-metagenome screening of BGCs. By analyzing 85,203 genomes and 9,428 metagenomes, new insights have been obtained about the diversity of BGCs on genomes from the majority of bacterial and archaeal lineages. This is exemplified by the discovery of the profound enrichment of BGCs in microbes after important geological events. Both the Great Oxidation and Cambrian Explosion events led to a surge in BGC diversity and abundance, particularly in polyketides. These findings suggest that microorganisms could adapt to the changing environment by evolving BGC to produce specific secondary metabolites. In summary, BGC-Prophet enables accurate and fast detection of BGCs on a large scale, holds great promise for expanding BGC knowledge, and sheds light on the evolutionary patterns of BGCs for possible applications in synthetic biology.

## Results

### BGC-Prophet model establishment and assessment strategy

BGC consists of a cluster of functionally related colocalized genes that can be regarded as sentences, and BGC prediction could be regarded as a problem of text classification in the field of natural language processing. Currently, many language models have been proposed and used to solve the problem of text classification, such as long short-term memory (LSTM) and bidirectional encoder representations from transformers (BERT). The original BERT proposed a revolutionary technique that generates generic knowledge of language by pretraining and then transfers the knowledge to downstream tasks of different configurations using fine-tuning [33, 34]. Following BERT’s mentality and paradigm, we developed a BGC language processing neural network model, BGC-Prophet, which captures location-dependent relationships between biosynthetic genes by being trained on thousands of microbial genomes and assigns gene types or product classes by simply plugging in two classifiers and fine-tuning the parameters supervised by a reference dataset (**Figure 1A-C**). Training on thousands of microbial genomes enables the model to learn the general syntax of genes, that is, gene location dependencies, which helps to improve generalizability and avoid overfitting. Fine-tuning ensures that the output embedding for each gene encodes context information that is more relevant to the biosynthetic functional profiles.

**Figure 1.**
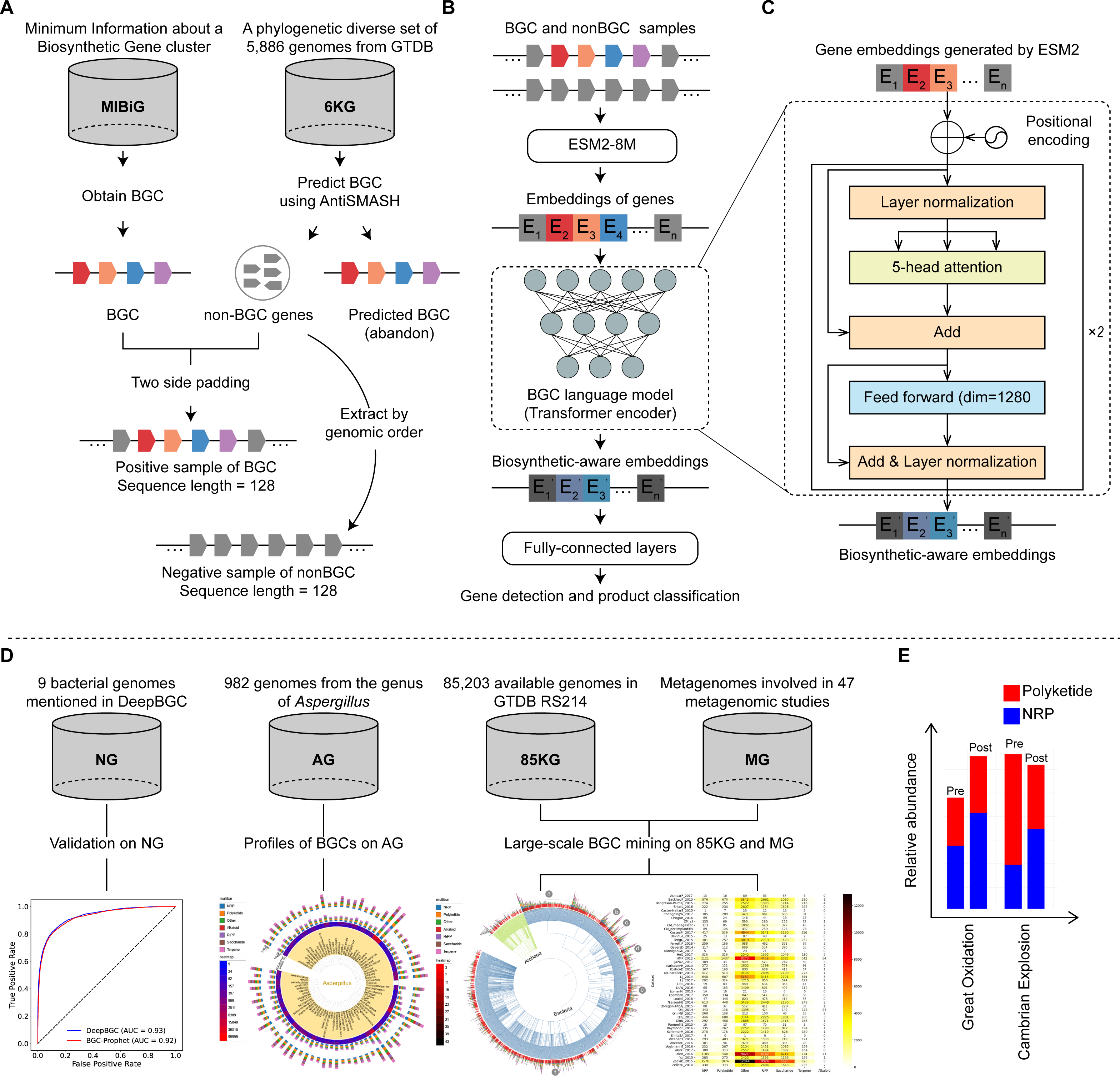
The workflow of our study. **A.** Generation of positive and negative samples. To train the BGC-Prophet language model, we curated a training dataset of 12,510 positive and 20,000 negative samples, and each sample was a cluster of 128 genes. MIBiG, minimum information about a biosynthesis-related gene cluster; 6KG, a phylogenetic diverse set of 5,886 genomes from the GTDB database. **B.** BGC-Prophet pipeline for BGC gene detection and product classification tasks. **C.** The architecture of the Transformer encoder used in BGC-Prophet. The prelayer normalization is used to accelerate the convergence of the model. The positional encoding adopts classical sine-cosine position coding, which does not require additional training and captures relative positional relationships between genes effectively. **D.** Several datasets used in this study for various purposes. NG, nine genomes that were examined in previous studies, such as ClusterFinder and DeepBGC; AG, 982 genomes from the genus *Aspergillus*; 85KG, 85,203 available genomes in GTDB RS214; MG, 9,428 metagenomics samples involved in 47 studies. Details of these datasets are available in **Methods**. **E**. The bar diagram shows the enrichment of BGCs in microbes after important geological events in Earth’s history.

BGC-Prophet has innovative designs to unleash its power in the BGC prediction task. First, BGC-Prophet uses genes as tokens to represent sentences (**Figure 1A**). Previous methods such as DeepBGC take Pfam domains as tokens that effectively balance genetic information loss and computational complexity. However, Pfam relies on manual determination by experts to define the scope of each domain. Here, we choose genes as tokens, which are more natural and do not require additional operations. Second, BGC-Prophet uses the evolutionary scale modeling (ESM, a pretrained language model for proteins) method to generate the embedding of gene tokens [35] (**Figure 1B-C**). The resulting numerical vectors of genes encapsulated evolutionary signals and functional properties based on their sequences, allowing us to leverage contextual similarities between genes.

To train the language model, we curated a training dataset of 12,510 positive and 20,000 negative samples, each of which is a gene cluster containing 128 genes (**Figure 1A, Supplementary Table S1**). Considering that the longest BGC in MIBiG (Minimum Information about a Biosynthesis-related Gene cluster) consists of 115 genes and the number of non-BGC genes between BGCs in genomes, we set the maximum number of genes to 128 in a sample (**Supplementary Figure S1**). Details of the generation of positive and negative samples are provided in the **Methods** section.

BGC-prophet accepts a set of genes as input and predicts BGC location and category. The input of the BGC-Prophet model is a sequence of embeddings represented by 320-dimensional vectors generated by the evolutionary-scale modeling (ESM) method [35] (**Supplementary Figure S2**). The output of the BGC-Prophet model consists of two parts. The first part is a sequence of values ranging from 0 to 1 representing the prediction scores of individual genes to be part of a BGC. The second part is which of the seven categories (see **Methods**) the input gene clusters belong.

We clarified several experiments for the evaluation and application of BGC-Prophet in this study (**Figure 1D**). First, we evaluated the performance of BGC-Prophet on the NG dataset, which comprises nine genomes mentioned in previous studies (**Supplementary Table S2**) [23, 26]. Second, we compared the BGCs predicted by BGC-Prophet and antiSMASH on the AG dataset, which comprises 982 genomes from *Aspergillus*, a genus with great biosynthetic potential. Then, we attempted to discover new insights into the diversity and novelty of BGCs on the 85KG, which comprises 85,203 available bacterial and archaeal genomes in the genome taxonomy database (GTDB), and MG (9,428 metagenomic samples involved in 47 studies) datasets (**Supplementary Table S3**). We finally studied the enrichment pattern of BGCs in microbes after important geological events in life on earth (**Figure 1E**).

### Evaluation of context-aware representations of genes

The ESM method generated context-aware representations of genes, thereby serving as meaningful input features for the BGC prediction model. In this subsection, we investigate the effectiveness of using vector representations generated by the ESM method. To achieve this, we first used ESM-2 8M (version 2 with 8 million parameters) to generate the vectors for a set of genes. Then, we consolidated the numerous genes within each BGC into a singular representative BGC vector by averaging the vectors. We evaluated the representative vectors of all BGCs from the MIBiG database via t-distributed stochastic neighbor embedding (t-SNE) analysis. Subsequently, we reduced the dimensionality of the representative BGC vectors from 320 dimensions to 2 dimensions by the t-SNE method for improved visualization.

Different categories of BGCs demonstrate distinct patterns within the t-SNE dimensionality reduction plot (**Figure 2**). It is evident that the seven distinct categories of BGCs exhibit a concentrated distribution into three clusters (top right, bottom left, and bottom right). For instance, terpenes predominantly cluster in the bottom right, saccharides and RiPPs primarily cluster in the top right, and polyketides primarily cluster in the bottom left and bottom right. The remaining categories display a widespread distribution across all three clusters. The boxplot showed that the points of any two categories of BGCs exhibited clear separation on the scatter plot (**Figure 2A**), such as polyketide and terpene (t test, p < 0.001). We also analyzed the two-dimensional distribution of BGCs (positive sample) and non-BGCs (negative sample) in the training set. Despite the fact that there are areas in the graph that are exclusively occupied by BGCs (bottom right), there is substantial overlap between BGCs and non-BGCs on the scatter plot (**Figure 2B**), although their distributions are significantly different on both axes (t test, p < 0.001). Our findings demonstrated that the ESM method generated context-aware representations of genes and therefore helped the language model learn the location-dependent relationships between genes that distinguish between BGCs and non-BGCs.

**Figure 2.**
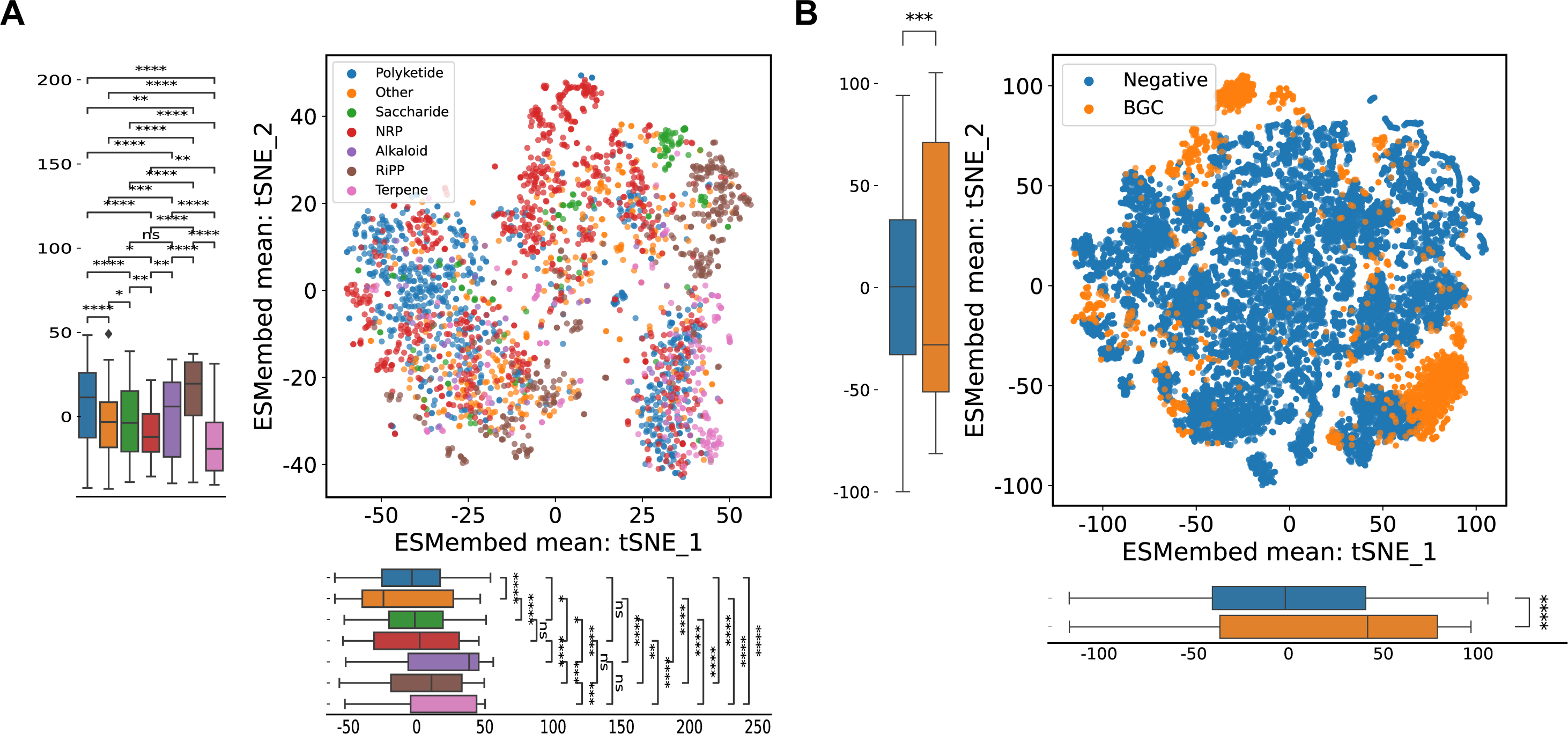
The distribution of ESM embeddings of genes. **A**. Average ESM embedding distribution for BGCs in the MIBiG database. The representative vectors of all genes within each BGC were averaged, and subsequently, a dimensionality reduction technique called t-SNE was applied to project the BGCs of all categories into a two-dimensional space. The resulting tSNE1 and tSNE2 values were then utilized to generate scatter and boxplot visualizations. The scatter plot depicts the spatial distribution of the average vectors representing the BGCs in the two-dimensional plane. Each BGC exhibits distinct distribution characteristics, and their separability is achieved through nonlinear means. Conversely, the boxplot graph displays the distribution patterns of the seven BGC categories along the tSNE1 and tSNE2 dimensions. It is evident that they predominantly occupy a specific region (-50,50), with only marginal discrepancies observed in terms of median values and distribution ranges. Importantly, the observed differences in distribution among the groups are statistically significant under a predetermined level of significance (*, p < 0.05; **, p < 0.01; ***, p < 0.001). NRP, non-ribosomal peptide; RiPP, ribosomally synthesized and post-translationally modified peptide. **B**. Average ESM embedding distribution for BGCs and non-BGCs. Using the t-SNE analysis as before, both the BGCs and non-BGCs were subjected to dimensionality reduction, resulting in scatter and boxplot visualizations. The scatter plot reveals that the non-BGCs are widely distributed, while the BGCs are primarily concentrated in the upper-left and lower-right corners of the plot. This indicates a clear distinction in the distribution patterns between BGCs and non-BGCs. On the other hand, the boxplot graph demonstrates that BGCs tend to be located at the edges of the plot, whereas non-BGCs exhibit a preference for the central region. This significant difference in distribution highlights the contrasting characteristics between BGCs and non-BGCs. We plotted separate boxplots for the horizontal and vertical axes and applied a t test to indicate that there were significant differences between pairwise comparisons of the samples at the given level of significance (*, p < 0.05; **, p < 0.01; ***, p < 0.001).

### Accurate and ultrahigh-throughput BGC prediction

To demonstrate the capabilities of our proposed framework, we assessed the performance of BGC-Prophet by evaluating its ability to (1) accurately locate BGCs throughout the bacterial genome (BGC gene detection) and (2) categorize them into their respective categories according to the types of their products (BGC product classification). Since DeepBGC is widely used by the community and is one of the best tools among existing BGC prediction tools, we choose DeepBGC as a representative and compare the performance of BGC-Prophet and DeepBGC.

BGC-Prophet has shown superior performance to DeepBGC in terms of accuracy. We initially evaluated the performance of BGC-Prophet and DeepBGC for BGC gene detection, and the results showed that the performances of BGC-Prophet and DeepBGC were comparable on the NG dataset (**Figure 3A**, **B**). Under the default threshold of 0.5, the BGC-Prophet model outperforms DeepBGC in metrics such as false positive rate and precision, while it lags behind in metrics such as false negative rate and recall (**Supplementary Figure S3**). However, in terms of the AUROC, BGC-Prophet achieved an overall AUROC of 91.9% with regard to locating BGCs throughout the bacterial genome, while DeepBGC achieved 93.1% (**Figure 3B**). We further examined the performance of both tools on individual genomes and found that BGC-Prophet outperformed DeepBGC in several cases (**Figure 3A**). Specifically, BGC-Prophet had a higher AUROC than DeepBGC on three of the nine genomes, and DeepBGC had a higher AUROC on the remaining six genomes (**Figure 3A**). BGC-Prophet achieved the highest AUROC of 96.0% on the genome GCA_000158815 (NCBI accession), while DeepBGC achieved the highest AUROC of 96.0% on the genome GCA_000154945 (NCBI accession). Subsequently, we evaluated the performance of BGC-Prophet and DeepBGC on BGC product classification. In this task, BGC-Prophet achieved an AUROC of 98.8% with regard to differentiating among the seven BGC categories, while DeepBGC achieved 91.3% (**Figure 3C**). This indicates that BGC-Prophet is better at accomplishing the BGC product classification task.

**Figure 3.**
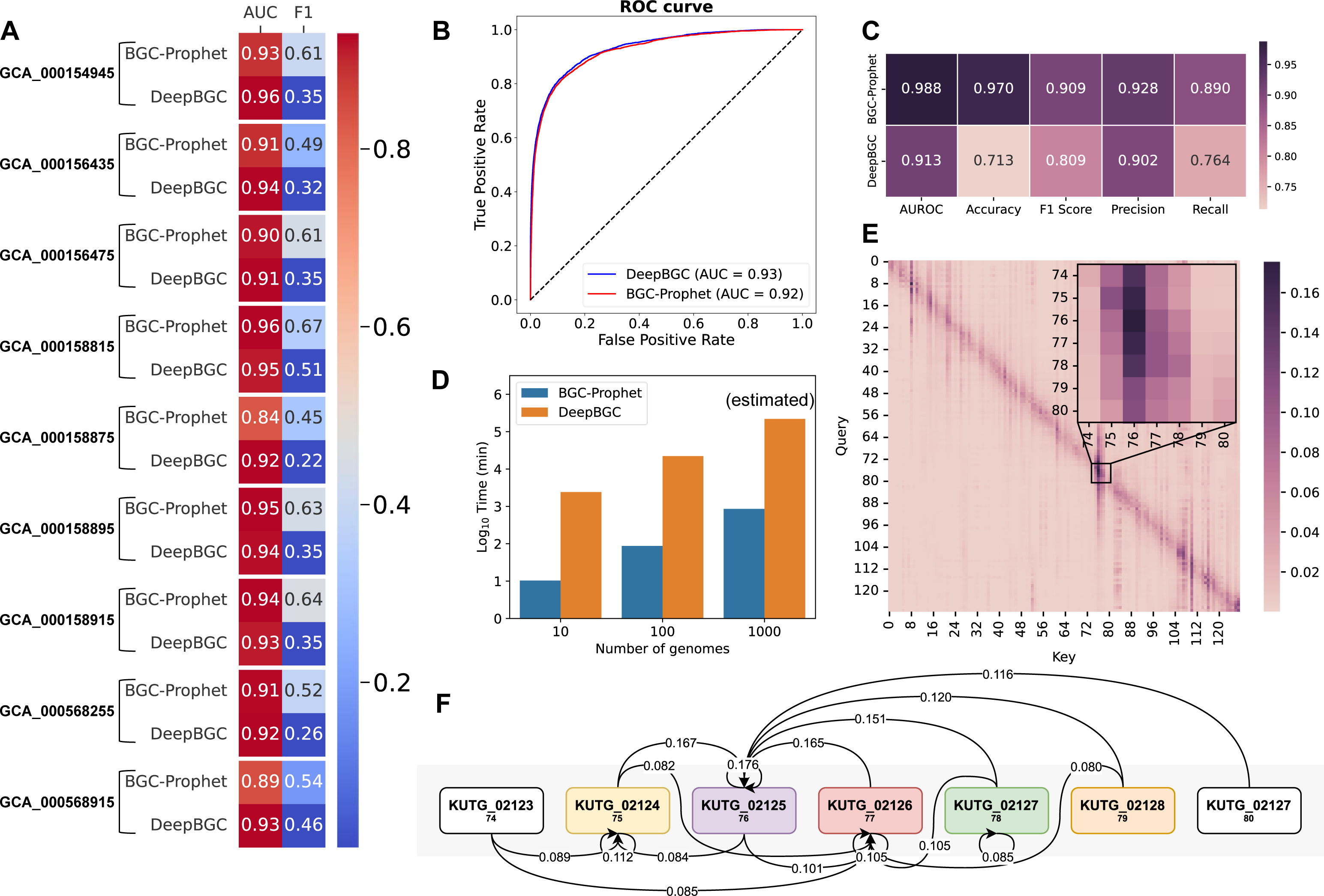
Evaluation of BGC-Prophet in different settings. **A**. The evaluation metrics reflecting the performance of BGC-Prophet and DeepBGC for BGC gene detection on the nine genomes that were examined in previous studies. All metrics except AUROC are evaluated under the default threshold of 0.5. **B**. The evaluation metrics reflect the performance of BGC-Prophet for BGC product classification, and the DeepBGC random forest classifier is retrained using the MIBiG database (version 3.0). **C**. The receiver operating characteristic curve reflecting the performance of BGC-Prophet. **D**. The running time of BGC-Prophet and DeepBGC on datasets with different numbers of genomes. **E.** The gene heatmap of a gene cluster (128 timesteps) during a single prediction process on the nine genomes. This heatmap illustrate the detection model’s first layer’s five heads (see **Methods**) average attention map during a single prediction process by BGC-Prophet on nine genomes. The vertical axis represents the Query in the self-attention mechanism, corresponding to the input gene embedding vectors, while the horizontal axis represents the Key. Horizontally, the large heatmap implies that determining whether a gene participates in forming a BGC requires considering information from multiple positions. Vertically, the vertical dark purple lines in whole genes heatmap represent a gene influencing the formation of a BGC by multiple genes. Darker colors along the diagonal in heatmap suggest that determining whether a BGC is formed primarily relies on the information embedded in its own vector, indicating the critical role played by the embedding vector’s information. The zoomed-in heatmap demonstrate the attention relationships between the BGC and surrounding genes. Genes 75-79 are annotated as BGC genes. **F.** The schematic diagram of attention applied between BGC genes and the genes at both ends, only colored genes belong to this predicted BGC. Panel **F** provides a schematic explanation of the magnified section in panel **E**, only attention scores between genes that exceeded 0.08 are shown as lines. The gene 76 (KUTG_02125), which encodes a non-ribosomal peptide synthetase, receiving the highest attention scores from other BGC genes. This suggests that annotation tasks need to consider information from this gene, possibly implying its conservativeness and centrality in this BGC.

BGC-Prophet uses a more efficient ESM method to generate vector representations of genes, avoiding the time-consuming sequence alignment (Pfams alignment), and improving the throughput of genomic data processing. For instance, when we extrapolate the number of genomes to 10 (randomly select and replicate genomes) for efficiency evaluation, DeepBGC needed an average of four hours per genome, whereas BGC-Prophet could process each genome in just one minute (**Figure 3D**). We emphasize that BGC-Prophet is the first UHT method that enables pan-phylogenetic screening and whole-metagenome screening of BGCs.

BGC-Prophet captures distant location-dependent relationships among biosynthetic genes. For example, we selected a BGC in the NG dataset and obtained its attention map during a single prediction process. The attention map shows the attention relationships between the BGC and surrounding genes (**Figure 3E**, **Supplementary Figure S4**). The gene 76 (KUTG_02125), which encodes a non-ribosomal peptide synthetase, receiving the highest attention scores from other BGC genes, possibly implying its conservativeness and centrality in this BGC (**Figure 3F**). Such examples are plentiful (**Supplementary Figure S4**), and the attention maps clearly show the language model can capture distant location-dependent relationships among biosynthetic genes.

### Comprehensive profiling of BGCs in 982 genomes from *Aspergillus*

BGC-Prophet predicts BGCs in a comprehensive manner and can predict more previously uncommented BGCs. Here, we utilized BGC-Prophet and antiSMASH to predict BGCs in genomes from *Aspergillus*, a genus with great biosynthetic potential and hundreds of genomes of this lineage. The results have shown that BGC-Prophet predicts a greater number of potential BGCs compared to antiSMASH, particularly in the terpene category (52,004 vs. 7,748, with 7,260 intersection BGCs). The predictions of BGCs in the NRP category by the two tools are nearly identical (27,603 vs. 27,100, with 26,278 intersection BGCs). BGC-Prophet predicts a larger number of BGCs in the categories of terpene and polyketide (35,606 vs. 18,225, with 16,607 intersection BGCs). Moreover, the prediction of BGCs in the RiPPs category by both tools exhibited complementarity (27,155 vs. 8,082, with 1,401 intersection BGCs), enhancing the coverage of predicted BGCs. Furthermore, BGC-Prophet predicts additional BGCs in the categories of alkaloids and saccharides compared to antiSMASH. The results showed a notable discrepancy between the BGCs predicted by the two tools, suggesting that BGC-Prophet can predict potentially novel BGCs beyond those detected by antiSMASH. We then studied the distribution spectrum of the predicted BGCs by both BGC-Prophet and antiSMASH. The results showed that BGC-Prophet predicted BGCs almost three times as many as antiSMASH (167,375 vs. 59,037, **Figure 4A**), and most of them are potentially novel BGCs (**Figure 4B**, **C**). The prediction results of the two tools showed a clear linear correlation (**Supplementary Figure S5**, r = 0.91, p < 0.001), indicating that the BGCs predicted by BGC-Prophet have no preference for specific species. Overall, we demonstrate that BGC-Prophet predicts BGC in a more comprehensive manner and can predict more previously unannotated BGCs.

**Figure 4.**
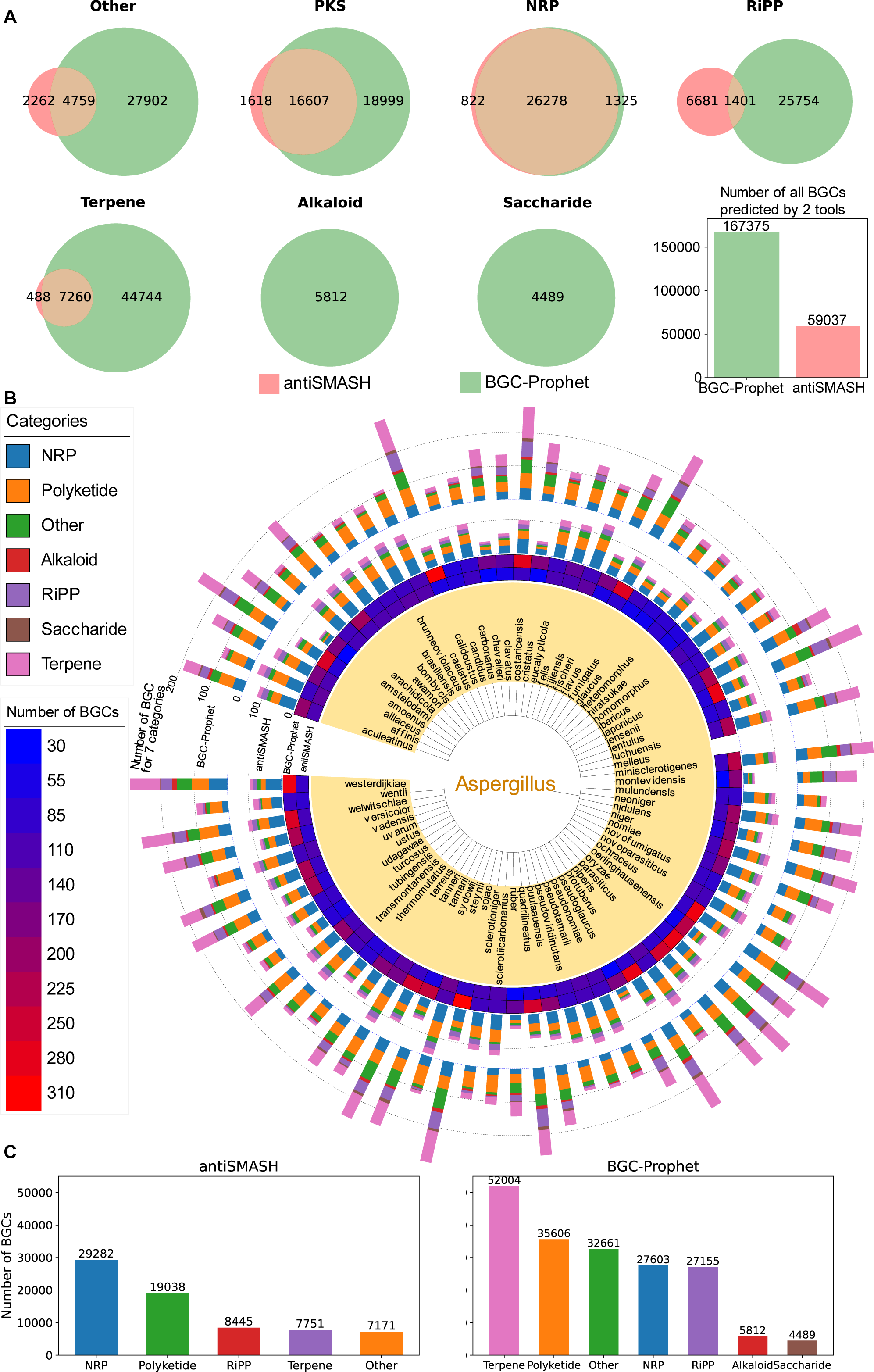
The predicted BGCs on the *Aspergillus* genomes dataset by BGC-Prophet and antiSMASH. **A**. The number of BGCs predicted by BGC-Prophet (green) and antiSMASH (red) for seven categories. The bar plot shows the total number of BGCs predicted by the two tools. Based on the assumption that if two prediction tools identify BGCs with identical genes, they are considered to have predicted the same BGC, the prediction results for all seven BGCs can be visualized using a Venn diagram for each category. It is important to note that antiSMASH does not predict BGCs belonging to the alkaloid and saccharide categories. The BGC-Prophet predictions are based on the default threshold of 0.5. **B**. The distribution of BGCs in the genomes of the *Aspergillus* genus. Within the central core, the encompassed area represents the entirety of *Aspergillus* species (a total of 76 species). The meaning of each circle from the inside out: first circle, the total number of BGC predicted by antiSMASH; second circle, the total number of BGC predicted by BGC-Prophet, third circle, the number of each category of BGC predicted by antiSMASH, fourth circle, the number of each category of BGC predicted by BGC-Prophet. Taking into consideration the presence of multiple subspecies genomes within a species, the number of predicted BGCs per species is averaged. **C**. A bar chart depicting the total number of different types of BGCs predicted by antiSMASH and BGC-Prophet reveals that BGC-Prophet predicts a significantly higher number of BGCs across various categories compared to antiSMASH.

### Comprehensive profiling of BGCs on 85,203 microbial genomes from the majority of bacterial and archaeal lineages

With BGC-Prophet, new insights have been obtained about the diversity of BGCs on genomes from the majority of bacterial and archaeal lineages. We used BGC-Prophet to investigate the profile of BGCs on 85,203 microbial genomes from the majority of bacterial and archaeal lineages. Among these genomes, 41,599 were found to contain BGCs, resulting in the identification of a total of 119,305 BGCs. We first performed an analysis to determine the proportions of different categories of BGC (**Figure 5A**). When we mapped BGCs to the species (**Figure 5B**), the three most widely distributed BGC categories were polyketide (34%), NRP (33%), and RiPP (24%), and the three most abundant categories were NRP (33%), polyketide (28%), and RiPP (27%). Conversely, the alkaloid category exhibited the narrowest distribution (2% of the total species) and the lowest abundance (1% of the total number, **Figure 5B**). In comparison, the three most abundant categories in the MIBiG database were polyketide (41%), NRP (34%), and RiPP (13%) [4]. Moreover, BGC-Prophet identified a significantly greater number of BGCs classified as the “other” category (increasing from 324 to 32,233 and from 13% to 24%), indicating its enhanced capability in mining potentially novel BGC categories. Our findings showed that BGC-Prophet identified several times more BGCs than MIBiG, with notable differences in the composition of BGCs (**Supplementary Table S4**).

**Figure 5.**
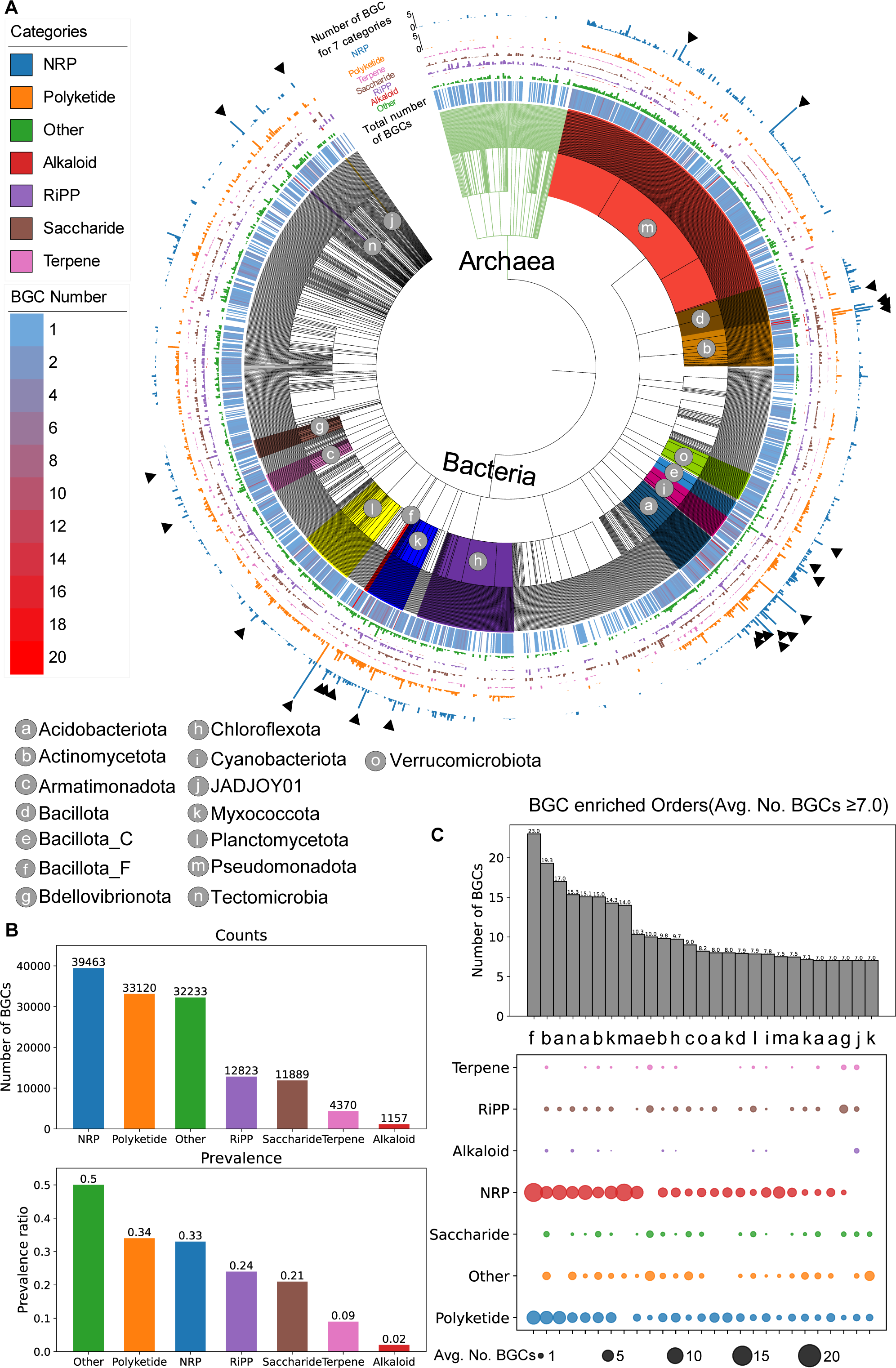
Novelty and phylogenomic distribution of microbial biosynthetic potential in different branches of the evolutionary tree of life. **A**. The distribution of predicted BGCs on an evolutionary tree is shown. The evolutionary tree consists of 85,203 bacterial and archaeal genomes from the GTDB database. For visualization purposes, the tree is displayed at the taxonomic rank of order. The number of BGCs is averaged per genome within each order. From the innermost to the outermost layers, the central core represents the evolutionary tree structure, consisting of 148 archaeal and 1,624 bacterial orders. The first ring depicts a heatmap of the total number of BGCs, with most genomes having two or fewer BGCs, while a few genomes exhibit up to 20 BGCs, indicating significant biosynthetic potential. The second to eighth rings display the distribution of BGC numbers for Other, Alkaloid, Saccharide, Terpene, Polyketide, and NRP categories, respectively. **B**. The term “prevalence” refers to the proportion of genomes that contain a specific type of BGC out of all the genomes analyzed. The sum of the prevalence values for the seven categories may exceed 100% because there can be instances where a single BGC belongs to several BGC categories. **C**. The average number of BGCs and the distribution of different BGC categories within selected orders. Among the orders, the top 27 orders with the highest average number (> 7.0) of predicted BGCs are distributed across 15 different phyla. These 27 orders represent a diverse range of phyla and showcase varying levels of BGC diversity and distribution across different BGC categories.

The host distribution of BGC showed species-specific characteristics, exemplified by the Actinomycetota phylum having the highest predicted number of BGCs (39,252 in total), and the Pseudomonadota phylum exhibited the widest genomic coverage, with 12,637 genomes containing at least one BGC, encompassing a total of 29,675 BGCs (**Figure 5A**, **C**, **Supplementary Table S5**). At the rank of order, the top 27 orders with the highest average number of predicted BGCs (> 7.0) are distributed across 15 phyla, such as Actinobacteria and Acidobacteriota (**Figure 5C**), which were reported to have relatively high biosynthetic potential (**Supplementary Text S1**). We proceeded to analyze BGCs separately for archaea and bacteria. Out of all these species, we identified 1,762 and 117,543 BGCs from 1,079 archaeal genomes and 40,520 bacterial genomes, respectively. On average, archaeal genomes contained 1.63 BGCs per genome, while bacterial genomes contained 2.90 BGCs per genome. These results indicate a significantly lower abundance of BGCs in archaeal genomes compared to bacterial genomes (t test, p = 6.1e-29). The predominant BGC categories in archaea were saccharides (30%) and RiPP (24%), whereas in bacteria, they were 10% and 11%, respectively. The predominant BGC categories in bacteria were NRP (33%) and polyketide (28%), whereas in archaea, they were 8% and 1%, respectively. This observation may be attributed to the more ancient nature of archaea compared to bacteria, particularly in energy acquisition and metabolism. While bacteria rely mainly on aerobic respiration, archaea have adapted to survive in extreme environments by using alternative strategies such as sulfur reduction, denitrification, and nitrate reduction (**Supplementary Text S2**) [36, 37].

### Comprehensive profiling of BGCs in 9,428 metagenomic samples

BGC-Prophet is the first UHT method that enables whole-metagenome screening of BGCs. We used 9,428 metagenomic samples corresponding to 47 studies from the human microbial environment and performed species annotations and BGC predictions on these samples (details in **Methods**). A total of 160,814 bins were generated from these metagenomic samples, of which 132,809 bins were successfully assigned to species, while 28,005 bins remained unclassified. Of the 9,428 metagenomic samples analyzed, a total of 8,255 were predicted to contain at least one BGC. The number of predicted BGCs was 248,229, distributed among 2,922 known species and unclassified species. The distribution of predicted BGCs from the human microbiome metagenomic dataset is shown in **Figure 6**. Consistent with the findings from the GTDB dataset, BGCs were significantly enriched in species belonging to Actinomycetota compared to species other than Actinomycetota (average of 8.30 BGCs per genome vs. 4.24 BGCs per genome, p = 1.06e-105). Additionally, archaeal species exhibited a higher number of BGCs compared to bacterial species (average of 9.00 BGCs vs. 4.88 BGCs, p = 0.0001). In terms of the abundance of BGCs for different categories, on average, there were 1.58 RiPPs, 1.36 saccharides, 1.12 NRPs, 1.10 polyketides, 1.07 terpenes, and 1.02 alkaloids.

**Figure 6.**
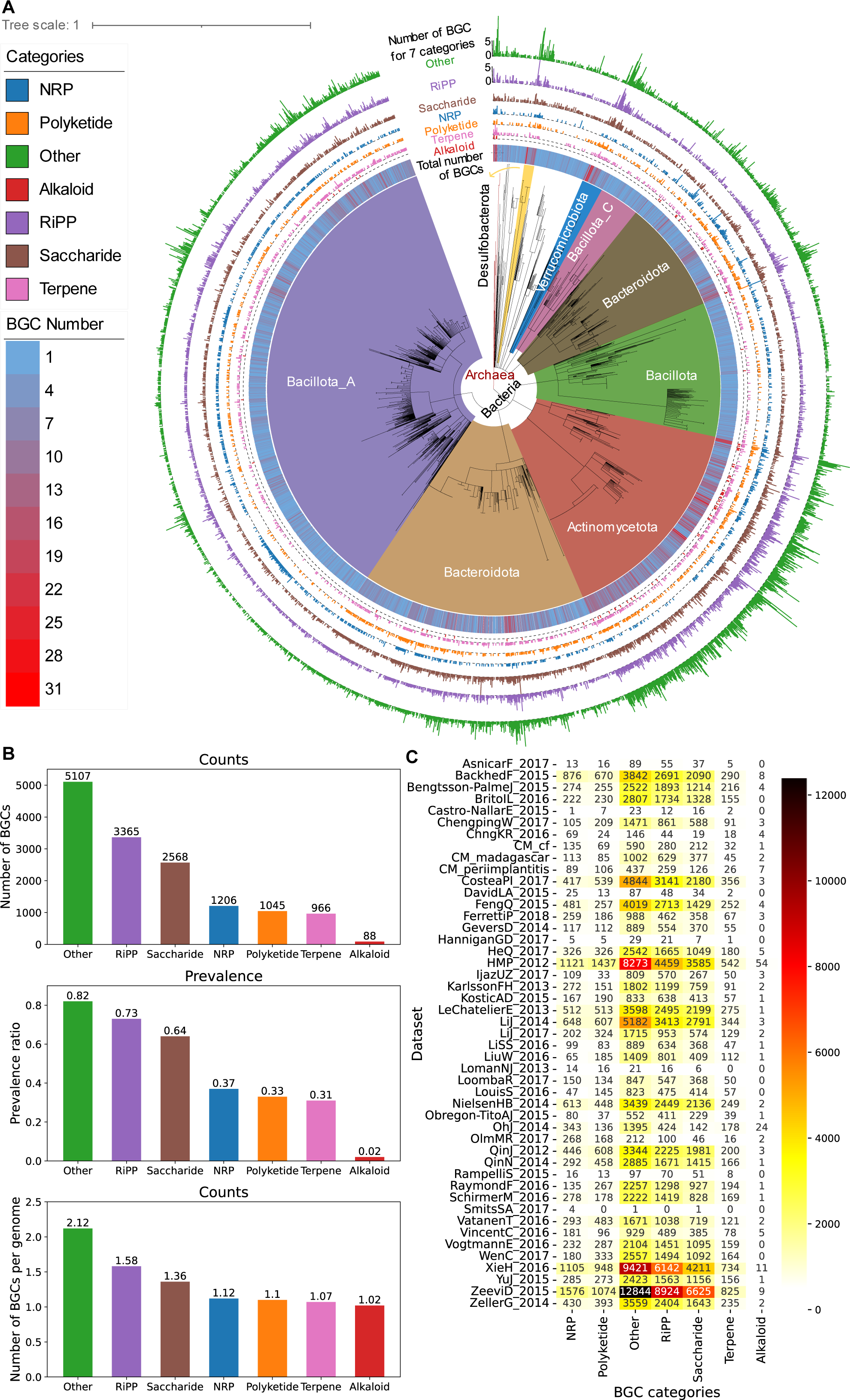
BGC-Prophet reveals the biosynthetic potential of the human microbiome. **A.** Distribution of predicted BGCs from the human microbiome metagenomic dataset (MG) on an evolutionary tree. BGCs were predicted using BGC-Prophet on the MG dataset, followed by species annotation. For the same species, the number of BGCs was averaged. From innermost to outermost, the central core represents the weighted evolutionary tree structure, composed of 13 archaeal species and 2,909 bacterial species, with different colored sectors representing several major phyla, such as Actinomycetota, Bacillota_A, and Bacteroidota. The first ring depicts the heatmap of the total number of BGCs, showing a clear enrichment in phyla such as Actinomycetota. The second to eighth rings display the distribution of BGCs belonging to the alkaloid, terpene, polyketide, NRP, saccharide, RiPP, and other categories, respectively. **B.** The bar plot from top to bottom shows the total counts, abundance, and average number of each type of BGC predicted. The order, from largest to smallest, is Other, RiPP, Saccharide, NRP, Polyketide, Terpene, and Alkaloid. The total count of BGCs is determined by the combination of abundance and the average number of a specific type of BGC per genome. The higher count of Other BGCs is likely due to shorter contig lengths, which result in fragmented BGC predictions that cannot be further classified. **C.** Heatmap showing the number of different types of predicted BGCs by BGC-Prophet in 47 metagenomic datasets. The predicted counts vary depending on the number of genomes, contig lengths, ecological niches, and biosynthetic capabilities of the respective datasets. For detailed information, please refer to **Supplementary Table S6**.

### The profound enrichment of BGC in microbes after important geological events

Large differences were observed in the distribution of BGCs among different species, particularly in light of the evolution over billions of years. To understand this phenomenon, we searched TimeTree [38] and identified two time points for the rapid growth of lineages, which corresponded to the Great Oxidation [39] and Cambrian Explosion [40] events (**Supplementary Figure S6**). After both of these events, we observed a surge in BGC diversity and abundance, possibly indicating the impact of the environment on BGC.

The Great Oxidation event occurred approximately 2.5 to 2.3 billion years ago, which was about the same time as the emergence of ribosomes [41, 42]. Prior to this time point, there was a shift in the evolution of certain bacterial genera, such as *Mesoaciditoga* [43], *Vampirovibrio* [44], and *Synechococcus* [44], which are categorized as the “pre” group. These genera comprise 56 out of 41,599 genomes analyzed. The remaining 2,215 genera evolved after this time point and are categorized as the “post” group. Statistical analysis revealed a significant increase in the average number of BGCs per genome from 2.5 to 4.5 between these two groups (t test, p = 0.024). The abundance of polyketide BGCs also showed a significant increase after the Great Oxidation event, with the average number of polyketides per genome rising from 1.09 to 2.81 (t test, p = 0.057). The possible reason is that polyketides are usually small compounds involved in oxidation reactions influenced by the increase in oxygen levels [45]. On the other hand, there were no significant differences in the average abundance of RiPPs (decreased from 1.29 to 1.25, t test, p = 0.807) and NRPs (increased from 1.0 to 3.16, t test, p = 0.242). The change in RiPP before and after the Great Oxidation event is not significant, which may be because the synthesis of RiPP primarily relies on the ribosomal pathway, involving dehydration and condensation reactions rather than oxidation [46]. On the other hand, although the number of NRP increases, the statistical significance is not substantial due to the limited data available before this event, which consists of only three cases [47].

The Cambrian Explosion event occurred approximately 542 to 520 million years ago, marked by rapid diversification of multicellular organisms [48]. Prior to this time point, 1,529 genera comprised 9,212 out of 41,599 genomes analyzed, which were categorized as the “pre” group. The remaining 589 genera evolved after this time point and were categorized as the “post” group. At this time point, there was a significant increase in the average number of BGCs per genome, with the “post” group having double the number compared to the “pre” group (6.07 vs. 2.95, t test, p = 4.89e-305). Further analysis of different categories of BGCs revealed significant differences in their average abundance before and after this time point. All categories of BGCs showed an increase in average abundance, with rapid increases observed in polyketides and NRPs. Polyketides encompass compounds such as erythromycin and tetracycline, while NRPs encompass cephalosporins, daptomycin, and vancomycin, among others. These compounds play crucial roles in defending against other bacteria and enhancing fitness in diverse environments [49, 50]. One possible explanation for this finding is that during the Ediacaran period, approximately 635-541 million years ago, Cyanobacteria began to appear, leading to a significant increase in oxygen production through photosynthesis, which resulted in heightened ocean oxygenation [51]. This amplified ocean and atmospheric oxygenation may have sped up the process of life evolution [52, 53]. It was during this time that multicellular organisms started to emerge [54]. On the one hand, multicellular organisms have always been hosts of microorganisms, and there is evidence to suggest that the genetic evolution of multicellular organisms occurred five times faster during the early Cambrian [55], leading to rapid life evolution in the oceans. On the other hand, the Earth’s ecological environment underwent alterations due to the activities of various species, generating numerous microenvironments [56]. These microenvironments provide a variety of environmental pressures for microbial selection, resulting in a surge in the biosynthetic potential of microorganisms and leading to the synthesis of diverse secondary metabolites that enable microorganisms to better adapt to different environments and compete for resources.

These findings highlight the evolutionary dynamics of BGCs on a large temporal scale and shed light on the impact of environmental changes on the diversity and abundance of specialized metabolites produced by microbes. Further research is needed to explore the functional roles and ecological significance of these BGCs in the context of bacterial evolution and their potential applications in various fields, including medicine and biotechnology.

## Discussion

BGCs represent a promising source of natural products but are difficult to discover, express, and characterize. In this study, we developed BGC-Prophet to comprehensively identify known and predict potentially novel BGCs and their products. BGC-prophet is a supervised language processing neural network model that captures the location-dependent relationships between genes and learns biosynthetic-aware representations of BGCs based on their gene evolutionary patterns. These new properties make BGC-Prophet advantageous over previous methods, enabling it to accurately and quickly profile BGCs for a wide range of lineages from microbial genomes and metagenomes.

The novelty of this work is demonstrated in three contexts. First, BGC-Prophet utilizes the powerful language model of the transformer encoder, uses the context-aware representations of genes as input features, captures the distant location-dependent relationships among biosynthetic genes, learns biosynthetic-aware representations of BGCs based on their gene evolutionary patterns, and shows superior performance to existing tools such as DeepBGC. Specifically, BGC-Prophet achieved an AUROC of 91.9% with regard to locating BGCs throughout the genome and 98.8% with regard to differentiating among the seven BGC categories (**Figure 3A-C**). BGC-Prophet’s exceptional processing speed enables it to quickly analyze vast amounts of genomic data with high efficiency (**Figure 3D**), allowing for extensive profiling of BGCs in large-scale genomic and metagenomic data.

Second, BGC-Prophet is the first UHT method that enables pan-phylogenetic screening and whole-metagenome screening of BGCs and builds a comprehensive profile of BGCs on 85,203 genomes and 9,428 metagenomes from the majority of bacterial and archaeal lineages. We investigated the biosynthetic potential of the *Aspergillus* genomes and revealed numerous potentially novel BGCs missed by antiSMASH (**Figure 4**). Our examination of the BGC profile in the majority of bacterial and archaeal lineages revealed that BGC-Prophet allows for the detection of previously undiscovered BGCs, as well as reconstruction of a comprehensive picture of BGCs on genomes from the majority of bacterial and archaeal lineages (**Figure 5**).

Third, BGC-Prophet reveals the profound enrichment pattern of BGCs after important geological events, possibly indicating the impact of the environment on BGC. Specifically, the Great Oxidation event had a profound impact on microbial genomes, with a significant increase in the average number of BGCs per genome, particularly in polyketides. This suggests that polyketides may play an important role in oxidation reactions due to the increased oxygen levels during this time. The Cambrian Explosion event led to a significant increase in the average number of BGCs per genome, with polyketides and NRPs displaying the most pronounced growth. These findings suggest that microorganisms adapted to the rapidly changing environment by producing specific sets of secondary metabolites, including polyketides and NRPs.

BGC-Prophet is not without limitations. First, BGC-Prophet can only determine the category of BGC but cannot determine the actual small molecule as a product of BGC. It is rare to predict BGCs directly from small molecules, and more to predict BGCs by understanding small molecules and their associated microorganisms. Thus, it is possible to predict BGC in microbial genomes associated with small molecules and then use computational chemistry to screen and validate the BGC that matches the small molecules. Further work on establishing the connection between BGCs and small molecules is warranted. In addition, BGC-Prophet requires substantial, accurately annotated training data, while few current natural product databases offer comprehensive, well-curated data. Despite our improved performance, further work is needed to curate more diverse BGC databases that can be used to improve the training and validation of our model. Other possible improvements might include the discovery of new categories of BGCs, as well as the examination of the gain or loss of BGCs on a dynamic scale.

Taken together, the results of this work reveal unprecedented throughput in BGC discovery and annotation via language model. As the first UHT method for pan-phylogenetic screening and whole-metagenome screening of BGCs, BGC-Prophet builds a comprehensive profile of BGCs on genomes from the majority of bacterial and archaeal lineages, reveals the profound enrichment pattern of BGCs after important geological events. The BGC-Prophet could find a way to better understand BGC patterns and mechanisms, as well as in a variety of applications, including microecology protection and synthetic biology.

## Methods

### Datasets used in this study

We manually curated several datasets in this study, including MIBiG v3.1 (Minimum Information about a Biosynthetic Gene cluster [3]), 6KG (5886 genomes from the GTDB RS214 database [57]), NG (nine genomes used in ClusterFinder and DeepBGC [23, 26]), AG (982 genomes from the genus of *Aspergillus*), 85KG (85,203 available genomes in GTDB RS214 [57]), and MG (metagenomes from 47 metagenomic studies [58]). These datasets are used for a variety of purposes, with MIBiG and 6KG being used to build training and testing sets, NG and AG being used to validate and compare the performance of various methods, and 85KG and MG being used for large-scale genome mining of BGCs (**Supplementary Table S1** and **S2**).

#### The MIBiG dataset

The MIBiG dataset specification provides a robust community standard for annotations and metadata on BGCs and their molecular products, which contains 2,502 experimentally validated BGCs.

#### The 6KG dataset

The 6KG dataset comprises a set of phylogenetically diverse genomes that were manually curated in GTDB RS214, and it contains 5,886 genomes that spread across the bacterial evolutionary tree.

#### The NG dataset

The NG dataset comprises nine bacterial genomes that were examined in previous studies, including ClusterFinder and DeepBGC [23, 26]. These genomes involved a total of 291 BGCs, none of which were used for training.

#### The AG dataset

The AG dataset contains a total of 982 genomes from the genus *Aspergillus* in the NCBI genome database. We utilized BGC-Prophet and antiSMASH to mine BGCs in these genomes and generated a comparison map between the BGCs identified by antiSMASH and BGC-Prophet on the *Aspergillus* genomes.

#### The 85KG dataset

The 85KG dataset contains 85,203 available genomes in GTDB RS214. We utilized BGC-Prophet to mine BGCs in those genomes and built a comprehensive profile of BGCs on genomes from the majority of bacterial and archaeal lineages.

#### The MG dataset

The MG dataset contains metagenomes involved in 47 studies (**Supplementary Table S3**). These metagenomic data included 1,792,406,629 contigs from 9,428 metagenomic samples, of which 6,238,438 contigs with nucleotide sequence lengths greater than 20,000 were retained. All datasets are publicly available and shown in **Supplementary Tables S1-S3**.

#### Taxonomic classifications for metagenomes

We used 9428 metagenomic assemblies corresponding to 47 studies from the human microbial environment. These metagenomic assemblies were binned using MetaBAT2 (version 2.12.1), and a total of 160,814 bins (or MAGs) were obtained. Taxonomy annotation was then performed on the resulting bins using the Genome Taxonomy Database Toolkit (GTDB-Tk, version 2.3.2) with reference to GTDB release 214.0. A total of 160,814 bins were generated from 9,428 metagenomic samples. Among them, 132,809 bins were successfully assigned to species, while 28,005 bins remained unclassified and were designated Unclassified (5,875 bins), Unclassified Archaea (316 bins), or Unclassified Bacteria (21,814 bins), representing unknown species.

### Positive and negative sample generation

To train the language model of BGC-Prophet, we manually curated a training dataset of positive and negative samples. The MIBiG and 6KG datasets were used to build the positive and negative samples. Before generating positive and negative samples, we used antiSMASH (v6) to identify BGCs on a public reference set of 5886 microbial genomes (6KG dataset). For each reference genome, regions predicted to be part of a BGC were removed, and these pruned genomes without BGC-like regions served as the non-BGC gene library.

#### Positive sample generation

The positive samples are derived from the 2502 BGCs in the MIBiG dataset. For each BGC in the MIBiG dataset, we applied two-sided padding with non-BGC genes (as described in the previous paragraph) until the gene sequence length equaled 128. Considering that the longest BGC in MIBiG consists of 115 genes and the gap (*i.e.*, the average number of non-BGC genes) between BGCs in the genomes from the 6KG dataset (**Supplementary Figure S1**), we set the maximum gene sequence length of a positive sample to 128. We repeated the generation procedure five times for each BGC in the MIBiG dataset, resulting in 12,510 positive samples.

#### Negative sample generation

In the generation of a negative sample (non-BGC), a major challenge is to make non-BGC have a certain degree of similar genes with genes in BGCs but lack the semantic information preserved in BGCs (*i.e.*, the order of genes in BGC). To generate a single negative sample, a random region from the non-BGC gene library was selected, and a subregion containing 128 continuous genes was randomly picked from the selected region. In total, 20,000 negative samples were generated.

#### Labeling the samples

According to the MIBiG database, there are seven categories of BGCs, including alkaloids, non-ribosomal peptides (NRPs), polyketides, ribosomally synthesized and post-translationally modified peptides (RiPPs), saccharides, terpenes and others (**Supplementary Figure S1**). Notably, each BGC may have more than one category, so the prediction of BGC categories is a multi-label seven-category problem. For example, the positive sample derived from BGC with MIBiG accession of BGC0000356 was labeled with both the categories of Alkaloid and NRP. For all the negative samples, they are not labeled into any of the seven categories.

### BGC-Prophet implementation

#### Token of the BGC-Prophet model

In the field of natural language processing, the minimal semantic unit is called a “token”, which makes up sentences. BGC-Prophet is a language processing neural network that takes genes as tokens to represent BGC or non-BGC (sentence). Previous methods ClusterFinder and DeepBGC take Pfam domains as tokens that effectively balance genetic information loss and computational complexity. However, Pfam relies on manual determination by experts to define the scope of each domain, and the utilization of pHMMs for identifying conserved Pfam domains in sequences is computationally intensive. Therefore, a trade-off between the number of Pfam alignments and computational speed must be considered. The situation of multiple Pfam domains originating from the same gene requires the model to learn such relationships separately. Here, we choose genes as tokens, which are more natural and do not require additional operations.

#### Vector representation of token

Each gene present in the training and testing samples needs to be represented as a word embedding vector to serve as input for subsequent language models. We used the ESM-2 8M model (evolutionary scale modeling: pretrained language models for proteins, version 2 with 8 million parameters) to generate vector representations of genes. ESM is the SOTA general-purpose protein language model, which can be used to predict structure, function and other protein properties directly from individual sequences [35]. For every positive and negative sample, we applied the ESM-2 8M model to generate a vector representation of genes (embedding dimension of 320). The ESM-2 8M model generates embedding of genes and removes the dependence between acquiring vector representations of tokens and training language models. The vector representation of tokens generated by the ESM-2 8M model directly from individual sequences is more concise and breaks the limitations inherent in the training samples, thus providing a higher possibility of predicting unknown BGCs. All genes in the training data are inferred using the ESM-2 8M model, and the mean of the model’s last layer output is selected as the final word embedding for the sequence. This implies that our word vectors tend to represent higher-level information and can more effectively leverage GPU acceleration for computational processes.

#### Model architecture and configuration

Here, we proposed a BGC language processing neural network model, BGC-Prophet, to detect known and predict potentially novel BGCs from genome sequences. BGC-Prophet employs a language model (*i.e.*, transformer encoder) [33] for BGC identification and classification. The transformer encoder is a neural network model of a specific architecture that uses a multi-head self-attention mechanism to speed up training. The self-attention mechanism introduced in the transformer encoder makes it suitable for parallel computation and better than the RNN or LSTM in accuracy. In this study, PyTorch v2.0.0 was used to implement the transformer encoder structure of BGC-Prophet, which learns the representation of gene sequences for different downstream tasks.

In this study, the parameters of the transformer encoder are set as follows (**Supplementary Figure S2**). The input dimension is set to 320, which is equivalent to the dimensionality of the embedding generated by the ESM-2 8M model. Then, pre-layer normalization is used to accelerate the convergence of the model [59]. The positional encoding adopts classical sine-cosine position coding, which does not require additional training and captures relative positional relationships between genes effectively. The transformer encoder is configured with two 5-head self-attention layers and a dropout rate of 10%. The model is trained using the AdamW optimizer with a learning rate of 1e-2 and a batch size of 64. Given that the number of training epochs is not fixed, an early stopping strategy is employed, where the loss value of the verification set stops improving after 20 epochs without decreasing, and the model obtained from the epoch with the lowest loss value on the verification set is chosen as the final model.

#### BGC gene detection and product classification

We assigned two downstream tasks to BGC-Prophet. The first task is predicting the BGC gene loci of a given BGC sequence, and the second task is predicting the BGC category of a given region on a genome.

The first task for BGC-Prophet is predicting the BGC gene loci of a given gene sequence. Specifically, given the sequence to be predicted composed of multiple genes, determine whether each gene is part of a BGC according to the position relationship of all genes. There may be no correlation between the genes that make up the sequence to be predicted, while the gene tag sequence of each gene that makes up the BGC is related in order. Therefore, the task can be statistically modeled using a linear-chain conditional random field (linear-CRF) [60]. According to the linear-chain CRF algorithm, the input is the sequence to be predicted, and the output is the gene tag sequence. In this paper, the downstream neural network is set as a fully connected layer with 128 timesteps, and the weight of each timestep is shared, different from DeepBGC. After passing through the fully connected layer, the hidden state vector dimension of the transformer encoder is reduced from 320 to 128, then from 128 to 32, and finally to 1, which represents the probability score that a given gene is part of a BGC. The Gaussian Error Linear Unit (GELU) [61] is used as the activation function for each fully connected layer, and finally, the sigmoid activation function is applied. The final fully connected layer outputs a scalar between 0 and 1 that measures how confident the model is that the gene belongs to the BGC. The loss functions of the model in this paper are binary cross entropy, and the AdamW [62] optimization algorithm is used to make them converge.

The second task for BGC-Prophet is predicting the BGC category of a given region on a genome. According to the MIBiG database, there are seven categories of BGCs, including alkaloids, NRPs, polyketides, RiPPs, saccharides, terpenes, and others. We encode these categories using one-hot encoding and consider an all-zero vector to represent the non-BGC category. Notably, each BGC may have more than one category, so the prediction of the BGC category is a multi-label seven-category problem. The problem can be described as follows: Extracting the sequence of hidden state variables from the transformer encoder model *H* = (*h*_1_,*h*_2_,...,*h_n_*),*h_i_* ∊ ℝ^*k*^ and calculating the average hidden state 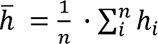. Transformer encoders allow input key padding masks to mask given specific timesteps, so this study uses gene tags as masks to prevent non-BGC genes from influencing the classification of BGC. The hidden states of the sequence are output as 7-dimensional vectors through a simple fully connected layer, and the sigmoid function is applied to output the confidence score of each label.

### Comparison methods

#### DeepBGC

DeepBGC is a novel deep learning and natural language processing strategy for improved identification of BGCs in bacterial genomes. DeepBGC employs a BiLSTM recurrent neural network. DeepBGC improves the detection of BGCs of known classes from bacterial genomes and harnesses great potential to detect novel classes of BGCs. In this study, we used DeepBGC for BGC gene detection and BGC product classification tasks and compared its performance to BGC-Prophet.

#### AntiSMASH

The antiSMASH (antibiotics & Secondary Metabolite Analysis Shell) is a comprehensive pipeline capable of identifying biosynthetic loci covering the whole range of known secondary metabolite compound classes. It employs a set of curated pHMMs to call biosynthesis-related gene families and a set of heuristics to designate a portion of a genome as a BGC. In this study, we applied antiSMASH and BGC-Prophet to 982 genomes from *Aspergillus* and evaluated the capability of BGC-Prophet to identify BGCs.

### Benchmark measures

Evaluation was based on five measuring metrics, including accuracy, precision, recall, F1-score, and AUROC. First, four parameters of the confusion matrix must be clarified: TP (true positive, actually BGC, and judged by the model as BGC), FN (false negative, actually BGC, but judged by the model as non-BGC), TN (true negative, the actual value is non-BGC, and judged by the model as non-BGC); and FP (false positive, the actual value is non-BGC, but judged by the model as BGC). We introduced several measures, including precision, recall, F1, true positive rate (TPR), and false positive rate (FPR). The definitions of these measures and formulas are as follows:

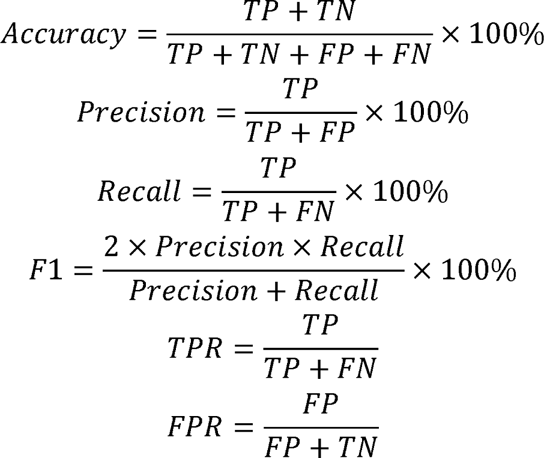

AUROC is the area under the receiver operating characteristic (ROC) curve, and ROC is the curve of TPR-FPR traversing different thresholds, which is also based on the confusion matrix.

### Statistical methods

UMAP (Uniform Manifold Approximation and Projection) and t-SNE (t-Distributed Stochastic Neighbor Embedding) dimensionality reduction techniques were applied to visualize and explore the high-dimensional gene vectors. To evaluate the differences between two BGC number groups, a t-test was performed. The t-test is a parametric statistical test that determines whether the means of two groups are significantly different from each other. It was used to compare the means of specific variables or features between the groups of interest. Pearson correlation coefficient was utilized to examine the linear relationship between the prediction results of the antiSMASH and BGC-Prophet. The Pearson correlation coefficient provides a measure of the strength and direction of the linear association between variables. It was employed to assess the correlation between different features or variables within the dataset.

## Declarations

## Ethics approval and consent to participate

Not applicable.

## Consent for publication

Not applicable.

## Competing interests

The authors declare that they have no competing interests.

## Authors’ contributions

K.N. and H.B. conceived of and proposed the idea and designed the study. Y.Z., Q.L., S.Y. and H.Z. performed the experiments and analyzed the data. Y.Z., Q.L., S.Y., K.N., and H.B. contributed to editing and proofreading the manuscript. All the authors have read and approved the final manuscript.

## Availability of data and materials

Data download links are provided in **Supplementary Table S3**. All source codes have been uploaded to the website at https://github.com/HUST-NingKang-Lab/BGCProphet. Detailed parameters of the software and package we used in this study are provided in **Supplementary Tables S1-S2.** All datasets and codes used in this study are publicly available.

## Funding

The National Key R&D Program of China (Grant Nos. 2021YFA0910500, SQ2023YFA1800082, 2018YFC0910502) and National Natural Science Foundation of China (Grant Nos. 32071465, 31871334, 31671374).

## Acknowledgments

We are grateful to Sugon (https://www.sugon.com/) for providing computational resources for this study.

---

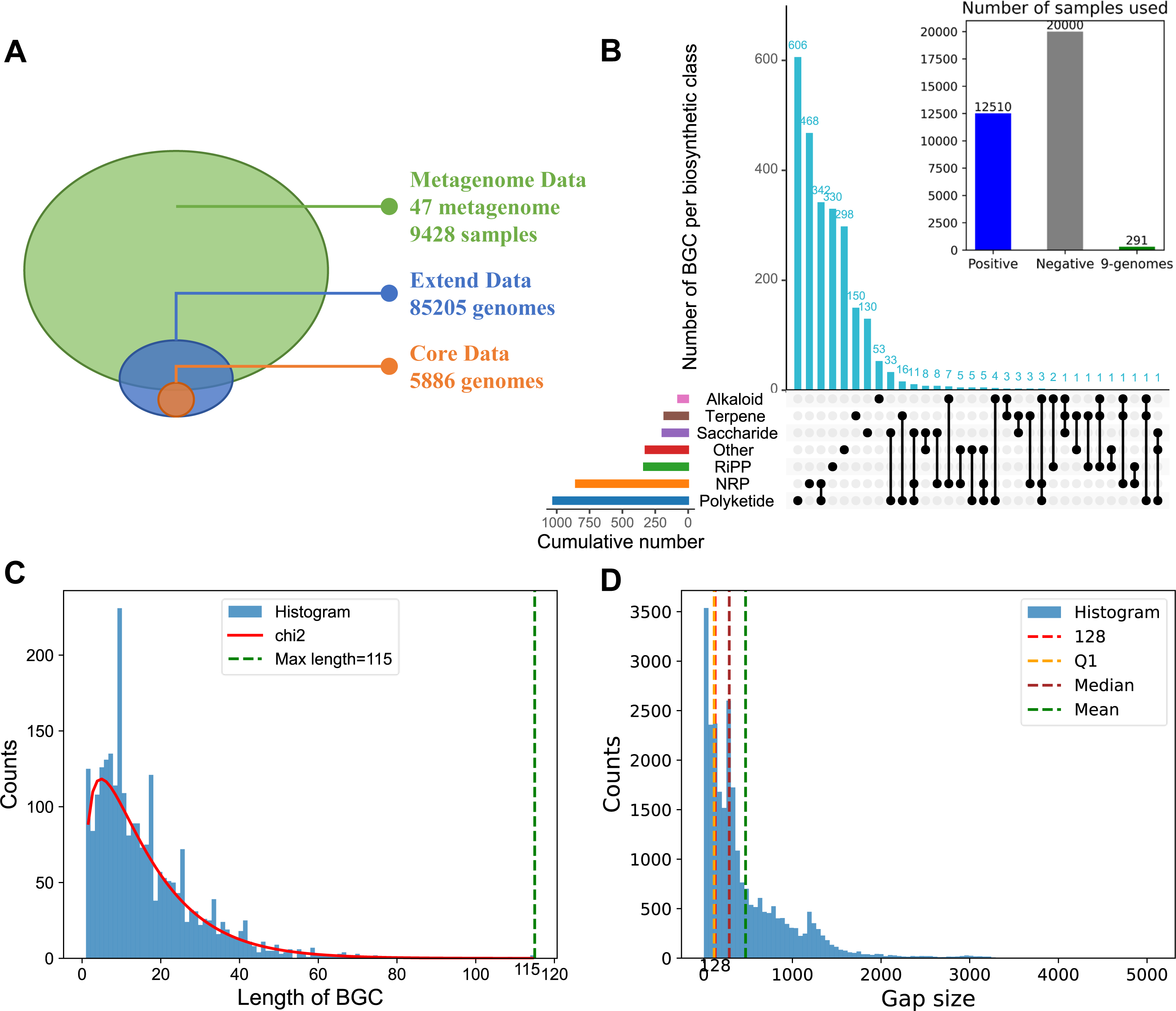

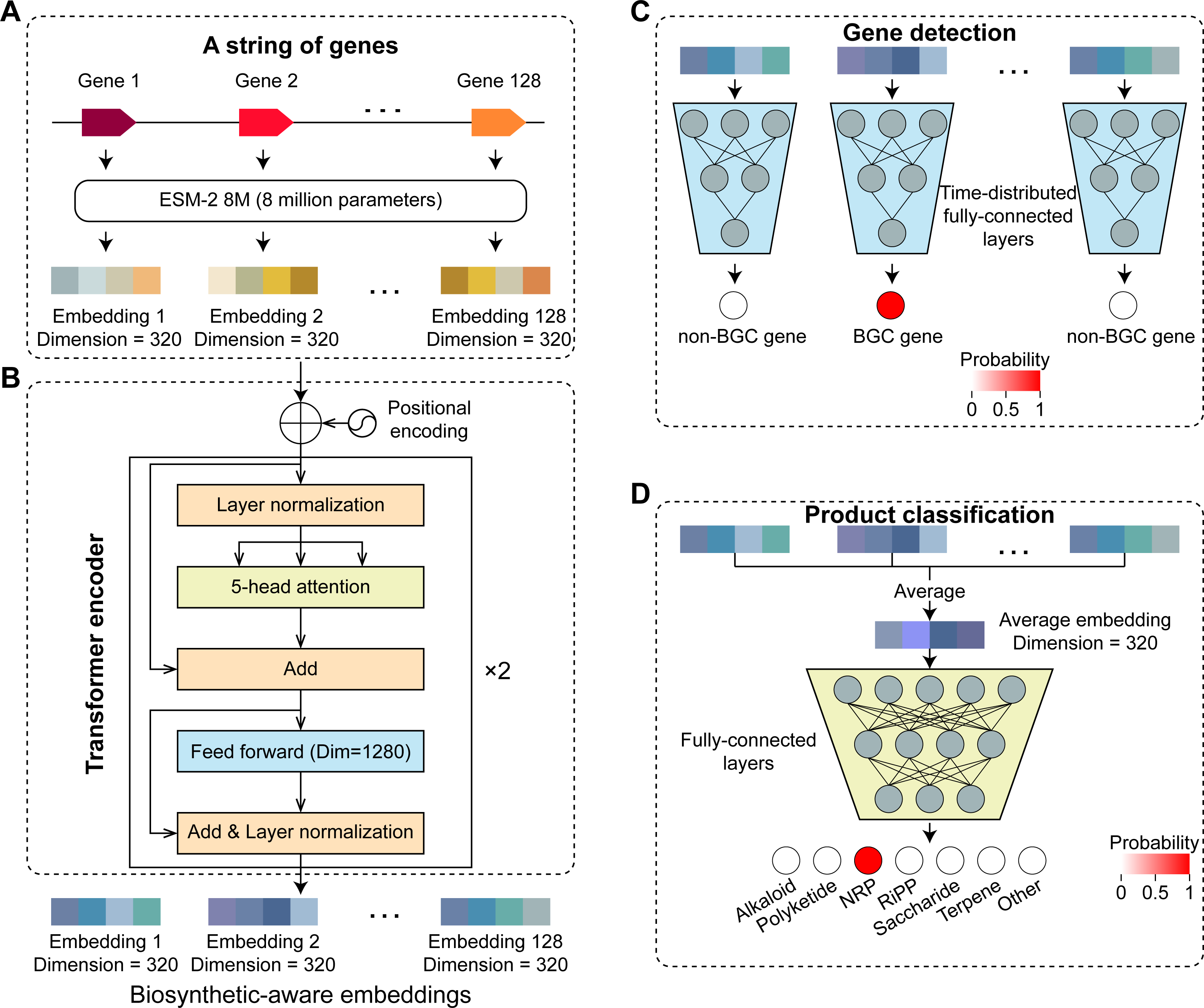

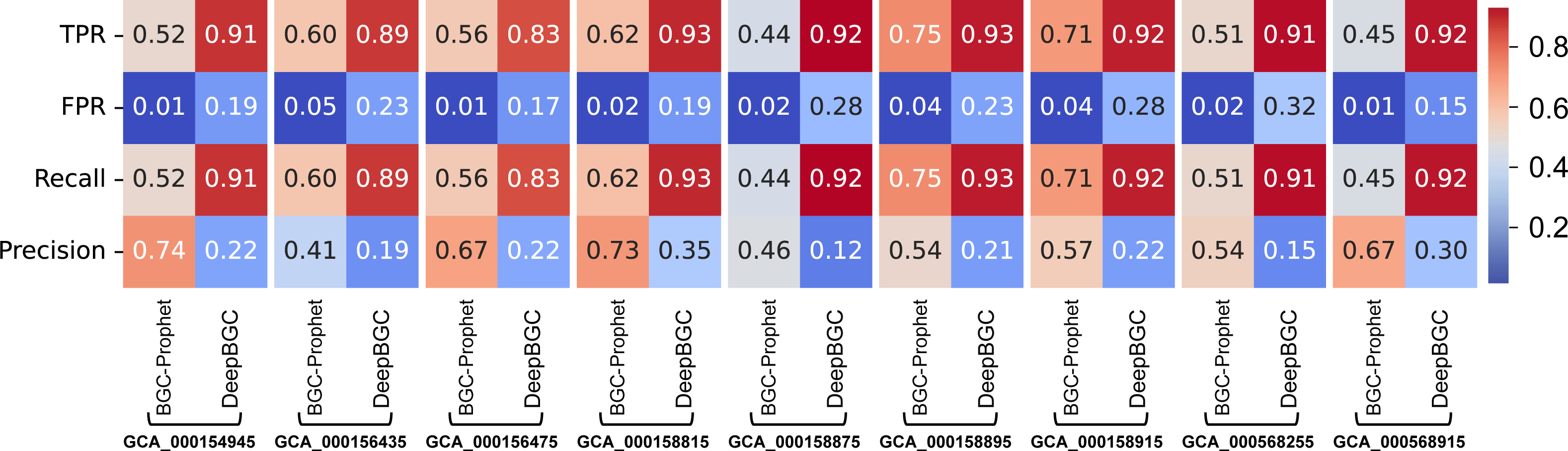

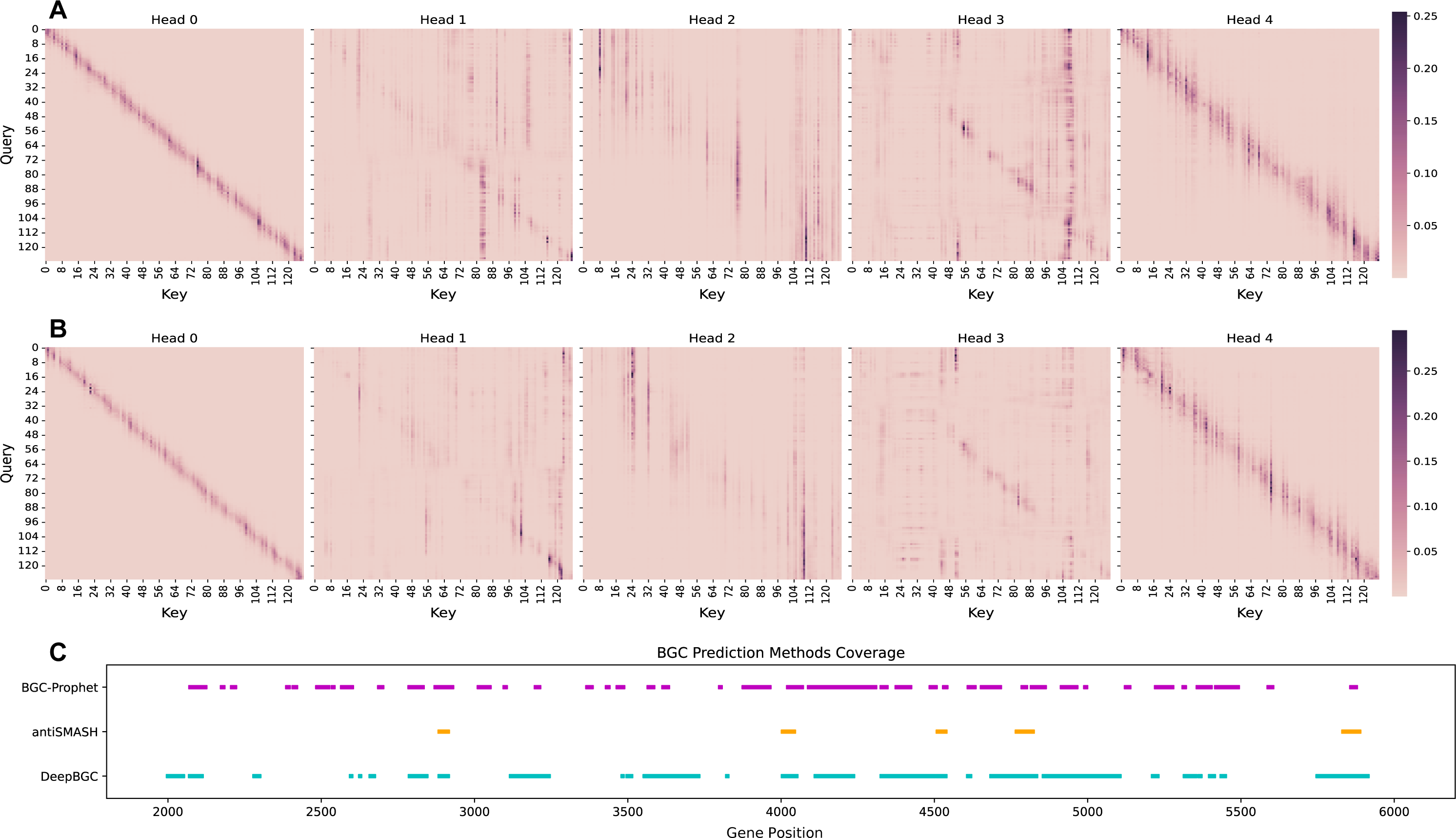

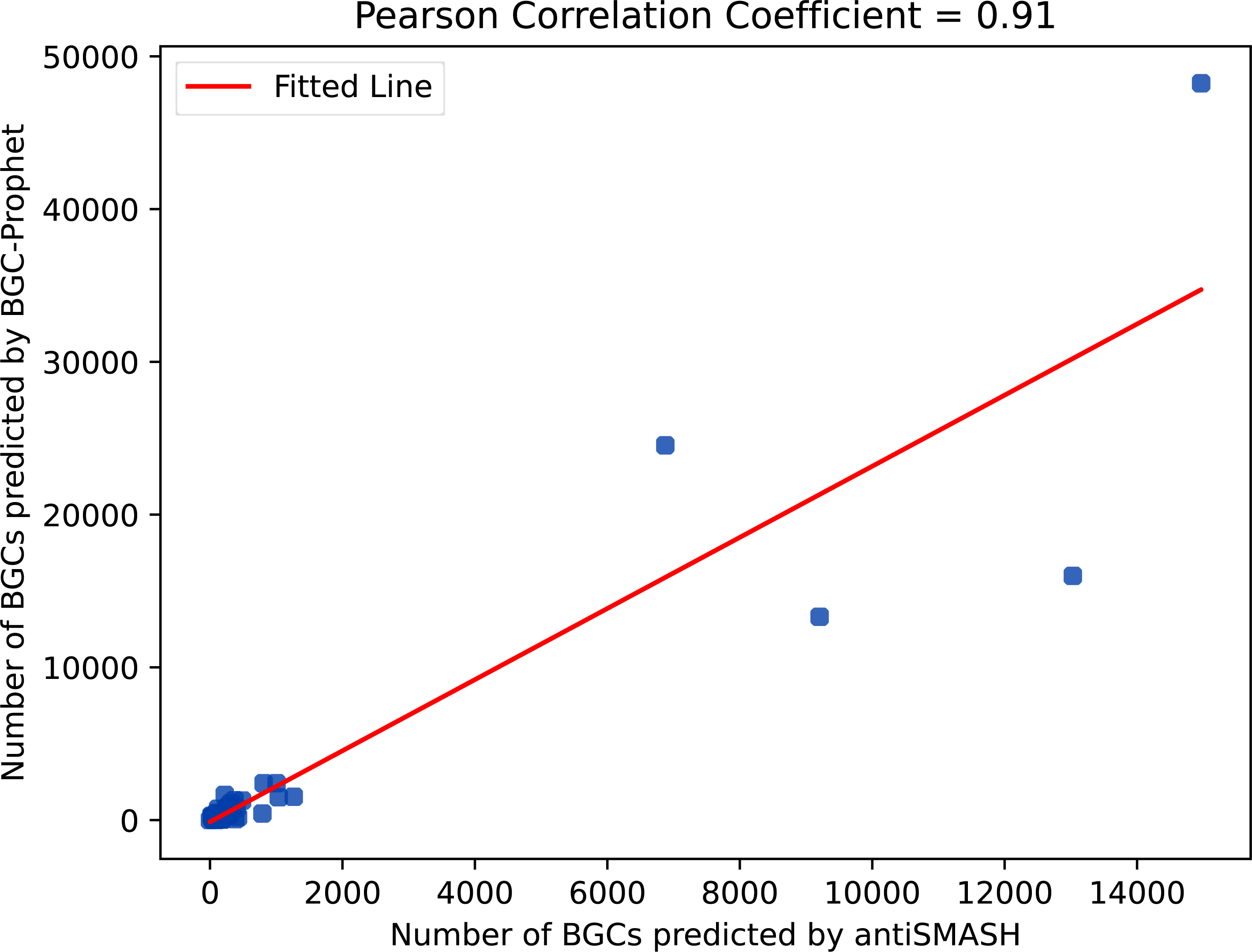

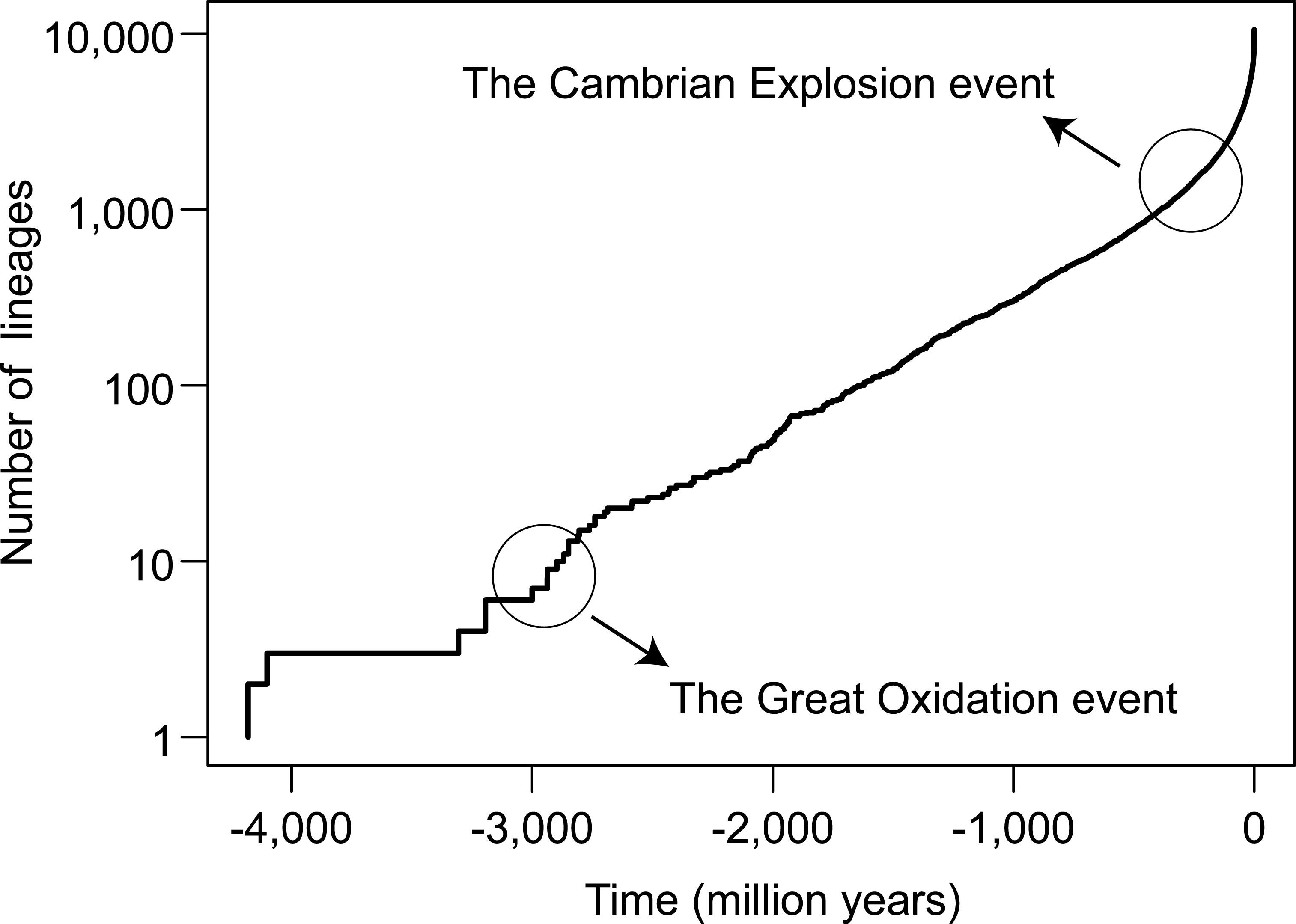

## Notes

https://doi.org/10.5281/zenodo.8104551

### Competing Interest Statement

The authors have declared no competing interest.

