## Supplementary Materials for "Deciphering the Biosynthetic Potential of Microbial Genomes Using a BGC Language Processing Neural Network Model"

**Running title**: *Lai Q et al / BGC-Prophet for BGC mining*

Number of Pages: 9

Number of Figures: 6

Number of Tables: 6

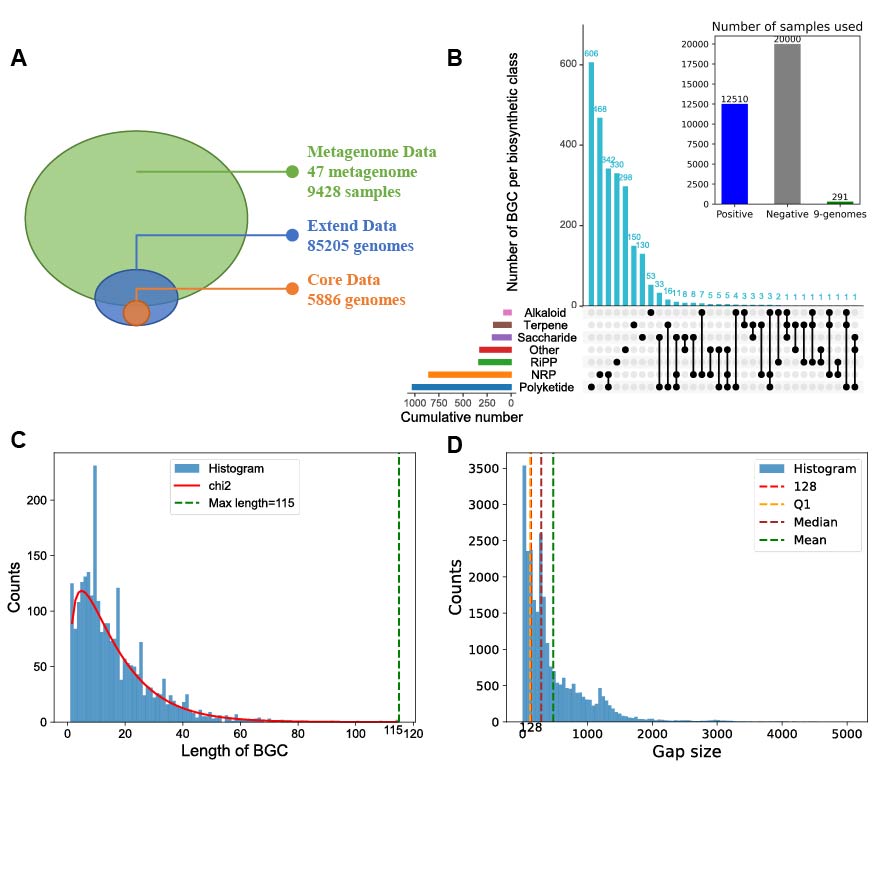
**Supplementary Figure S1. Dataset involved in this study and data statistics.** **A.** Description of the datasets used in this study. The core data contains 5,886 genomes used for building training and testing set, the extend data and metagenome data are used for large-scale BGC mining (**Supplementary Table S1**). **B.** The number of BGCs for each category in MIBiG database, and samples in the training and testing sets. **C.** Distribution of the number of genes in BGCs. **D.** Distribution of the number of non-BGC genes between two adjacent BGCs.

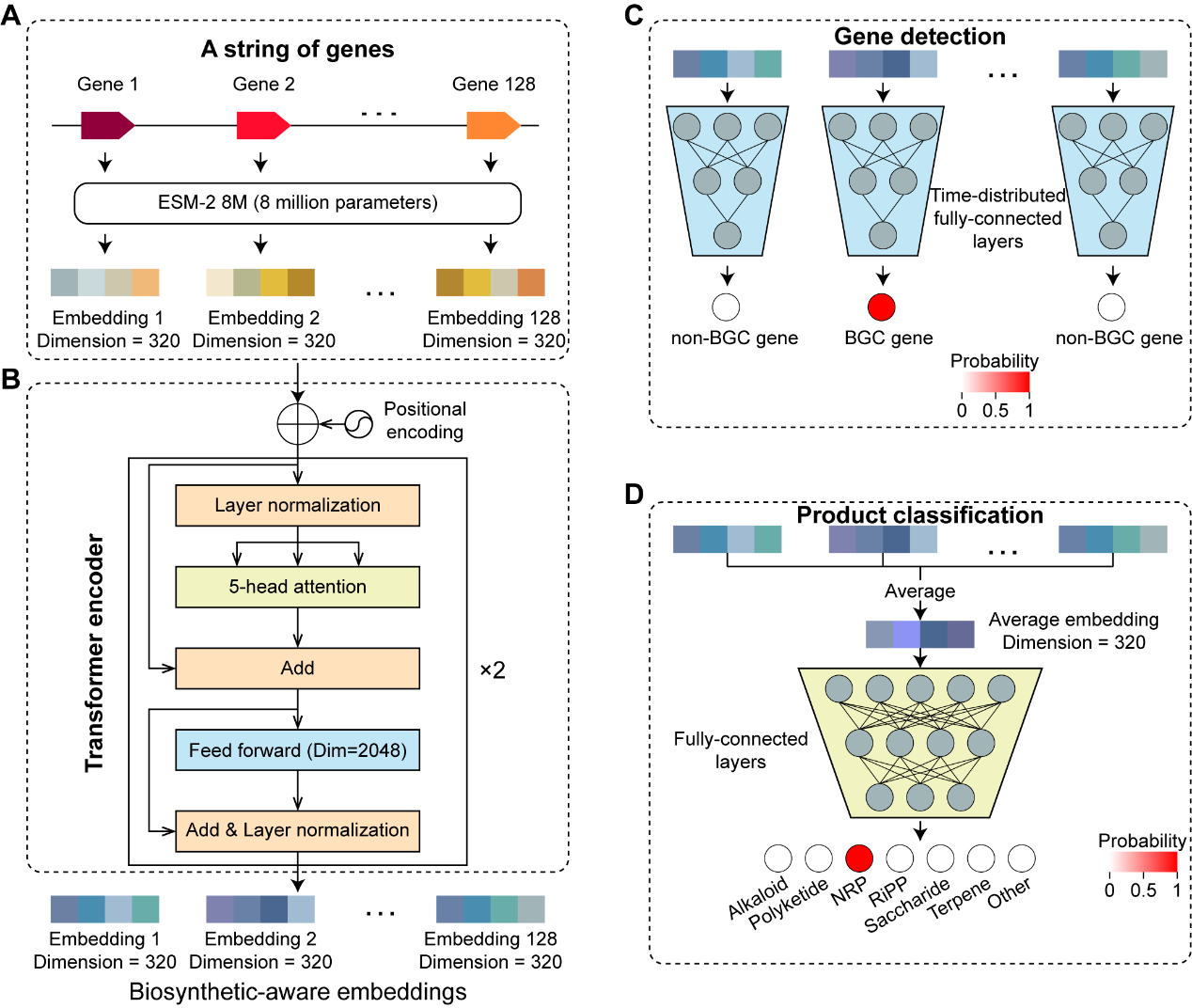

**Supplementary Figure S2. The architecture of the BGC language processing neural network model. A.** The input of model was a sequence of ESM-2 8M embeddings represented by 320-dimensional vectors. **B.** The BGC-Prophet model contains a transformer encoder module, which is used to learn representation of gene sequence. In this study, the parameters of transformer encoder module are set as follows. The input dimension is the same as the dimension (*i.e*., 320) of the embedding layer, and pre-layer normalization is used to accelerate convergence of model. Positional coding uses the classic sine-cosine position coding, which needn’t training and can capture relative position relationships. Transformer encoder was configured with two 5-head self-attention layers and a dropout of 10%. **C.** The classification module for gene identification was configured with time-distributed fully-connected layers and 128, 32, 1 output units. The output was a sequence of values between 0 and 1 representing the prediction scores of given genes to be part of a BGC. **D.** The classification module for product classification was configured with fully-connected layers and seven output units. The output was the seven values representing what BGC category the input sequence belongs to.

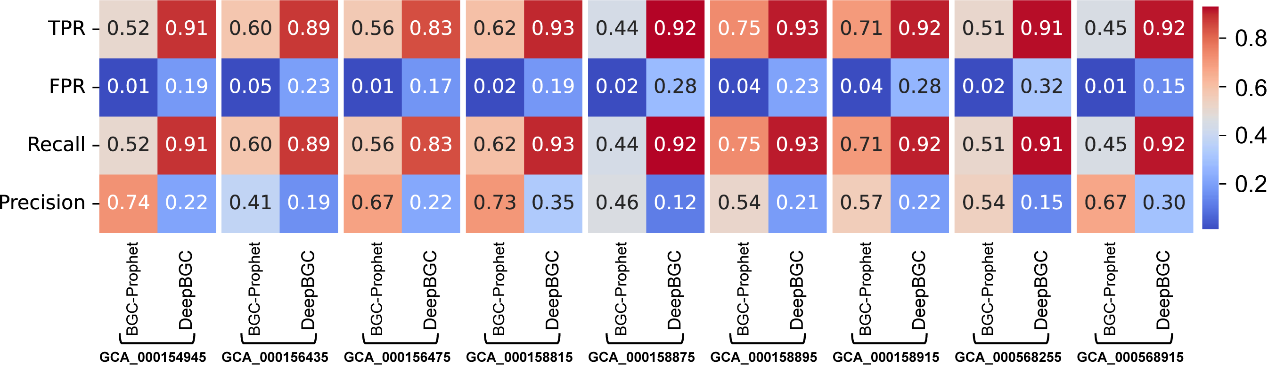

**Supplementary Figure S3. Evaluation of BGC-Prophet in different measurement metrics.** The evaluation metrics reflecting the performance of BGC-Prophet and DeepBGC for BGC gene identification on the NG dataset. The NG dataset comprises nine bacterial genomes used in previous studies, including ClusterFinder and DeepBGC. These genomes involved a total of 291 BGCs, none of which were used for training. All metrics are evaluated under the default threshold of 0.5. TPR, true positive rate; FPR, false positive rate.

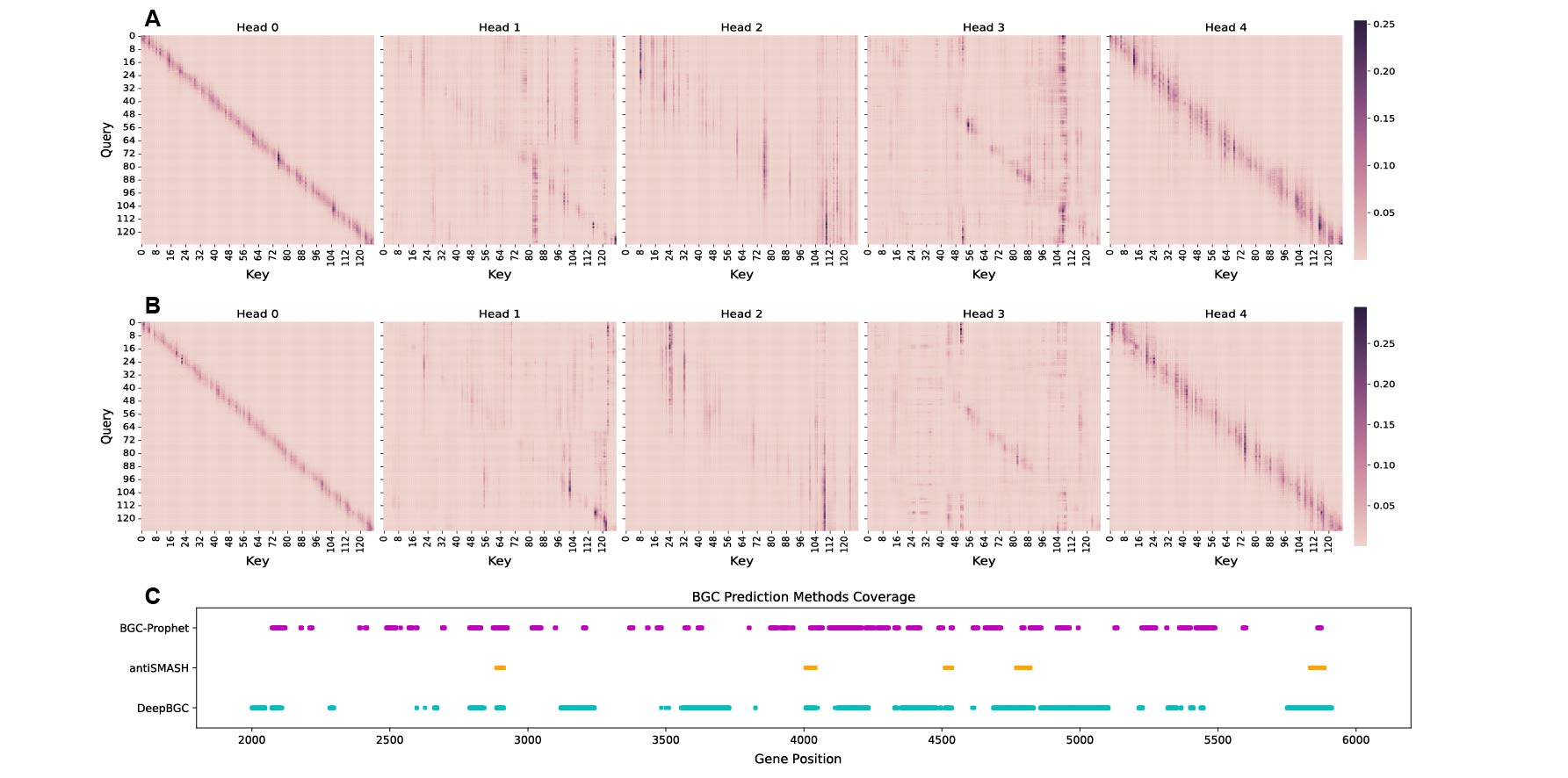

**Supplementary Figure S4. The attention map of BGCs and the coverage of different methods. A.** Five heads attention maps of detection on nine genomes. Based on the color intensity in the heatmap, it can be observed that different regions are attended to by the five attention heads (see **Methods**). Head 0 and 4 exhibit deeper colors along the diagonal, indicating that these attention heads focus on embedding vectors related to both the input and the surrounding genes. The distinction between Head 0 and 4 lies in the fact that Head 0 pays attention to fewer surrounding genes than Head 4. These two attention heads reflect a similar pattern to an RNN's attention to information from preceding and succeeding genes. Head 2 displays a deep purple vertical line at the position of gene 76, corresponding to the location of the BGC gene, suggesting that this attention head may be more concerned with the impact of the BGC gene on annotation. Head 1 and 3 also exhibit vertically aligned lines at different positions, indicating their potential focus on the impact of other genes on annotation. **B.** Five heads attention maps of detection from training sample. These five heatmaps represent the attention maps of the five attention heads in the first encoding layer of the well-trained detection model on the training samples, with positions 16-28 corresponding to the BGC gene. Similar to Figure A, Head 0 and 4 exhibit deeper colors along the diagonal, and Cases A and B further confirm the model's focus on both intrinsic and surrounding gene information. Head 2 also displays a deep purple vertical line at the position of the BGC gene (positions 16-28), while Head 1 and 3 show specific attention to other positions, although the precise patterns of attention in these cases are yet to be summarized. **C.** Three BGC prediction methods' coverage on Aspergillus genome. Three different methods were applied to the genome GCA_009687185.1, and genes predicted as BGC genes at each position were annotated on the graph. It is evident from the graph that antiSMASH identifies the fewest BGCs, while both DeepBGC and BGC-Prophet encompass the predictions made by antiSMASH. Additionally, there is an overlap between the predictions of DeepBGC and BGC-Prophet, suggesting their potential utility as cross-validation measures.

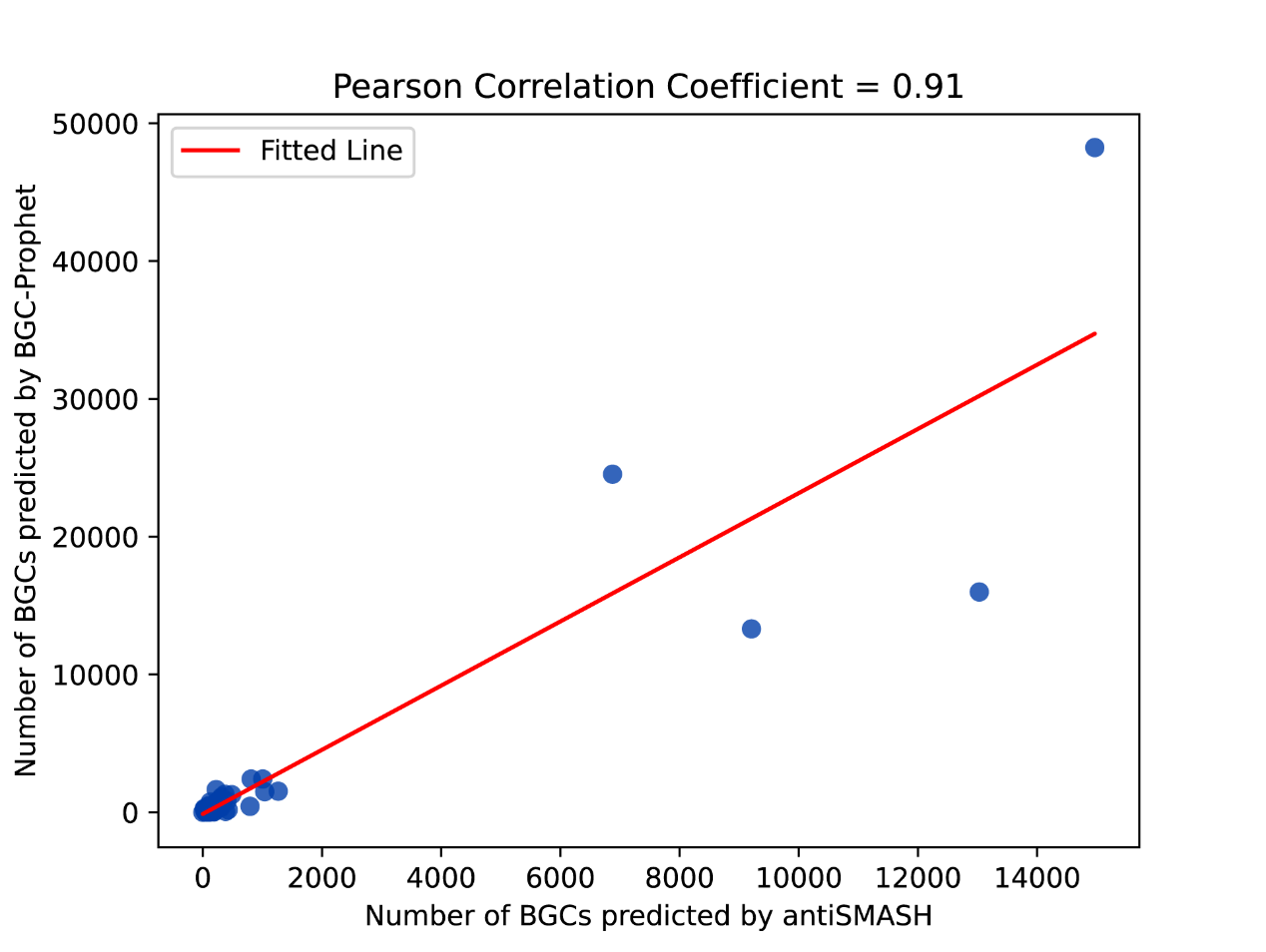

**Supplementary Figure S5. The number of BGCs predicted by BGC-Prophet and antiSMASH on the *Aspergillus* has a linear correlation (r =0.91, t-test, p <0.001).**

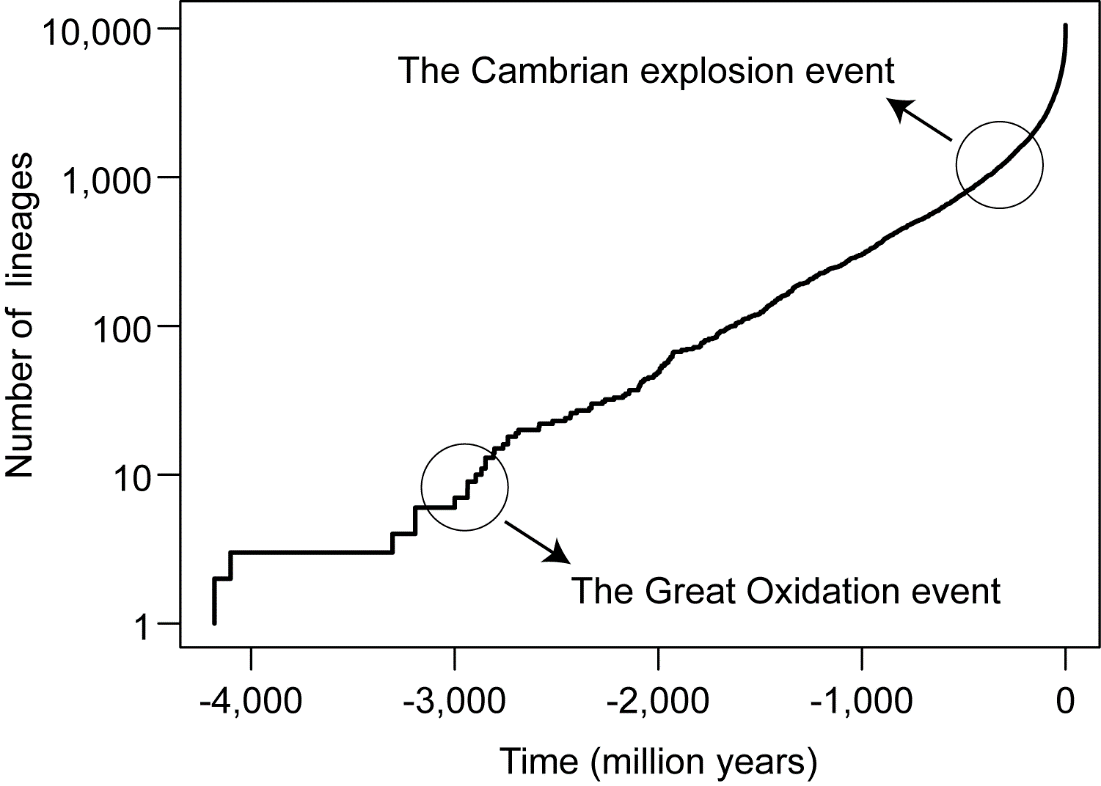

**Supplementary Figure S6. The increase in the number of bacterial and archaeal lineages over time.** We identified two time points for the rapid growth of lineages, which corresponded to the Great Oxidation and the Cambrian Explosion events in Earth history.

**Supplementary Table S1. Tools and databases used in this study.** Attach in Excel.

**Supplementary Table S2. Concepts used in this study.** Attach in Excel.

**Supplementary Table S3. Metagenomes used in this study.** Attach in Excel.

**Supplementary Table S4. The comparison of BGC composition between BGC-Prophet and the MIBiG database.** The BGCs are predicted by the BGC-Prophet on the 85KG dataset. Attach in Excel.

**Supplementary Table S5. The distribution of BGCs predicted by BGC-Prophet on the 85KG dataset at the rank of phylum.** Attach in Excel.

**Supplementary Table S6. The distribution of BGCs predicted by BGC-Prophet on the metagenomic dataset.** Attach in Excel.

**Supplementary Text S1. The Actinobacteria and Acidobacteriota have** **relatively high biosynthetic potential.**

Actinobacteria is one of the most morphologically diverse prokaryotes and is widely distributed in terrestrial and aquatic ecosystems [1]. In addition to utilizing unique biochemical pathways not found in other prokaryotes, they also synthesize many large molecules that are absent in other organisms, such as unique cell wall peptidoglycans [2]. Among them, the genus *Streptomyces* is one of the largest bacterial genera and is considered a producer of many biologically active metabolites that are useful in medicine for humans. These include antibiotics [3], antifungal agents [4], antiviral drugs [5], anticoagulants [6], immunomodulators, anticancer drugs [7], and enzyme inhibitors [8]. In agriculture, they are involved in the production of insecticides, herbicides, fungicides, and substances that promote plant and animal growth [9]. Antibiotics derived from Actinobacteria are particularly important in medicine and include aminoglycosides, anthracyclines, chloramphenicol, macrolides, tetracyclines, and others. Members of the Acidobacteriota can be found in various environments, including soil, decaying wood [10], hot springs, marine environments, caves, and metal-contaminated soils [11]. These bacteria are particularly abundant in soil habitats, accounting for 52% of the total bacterial community [12].

**Supplementary Text S2. The distinct mechanisms of energy acquisition and metabolism between archaea and bacteria.**

Archaea and bacteria exhibit distinct mechanisms for energy acquisition and metabolism. Bacteria primarily derive energy through aerobic respiration, whereas archaea are predominantly found in extreme environments such as concentrated saline waters, volcanic springs, anoxic depths of marine sediments, and the gastrointestinal tracts of animals [13]. In these habitats, archaea utilize unique strategies to acquire energy, including sulfur reduction, denitrification [14], and nitrate reduction [15].
